# Accumulation of Lipid Droplets in Microglia following Neonatal Brain Hypoxia-Ischemia

**DOI:** 10.64898/2026.07.18.739219

**Authors:** Fan Li, Yuqing Lei, Sihao Li, Gui Zhang, Yinuo Li, Beiyan Wu, Donna M Ferriero, Peipei Pan, Zhonghui Guan, Xiangning Jiang

## Abstract

**Background:** Hypoxic-ischemic encephalopathy (HIE) is a major cause of neonatal mortality and neurodevelopmental impairments. Following brain hypoxia-ischemia (HI), microglia face substantial metabolic stress; and upon phagocytosis, they become overloaded with lipids derived from engulfed dead neurons and myelin debris. It is unclear how microglia respond to and process the lipid cargo, and whether lipid accumulation may affect microglia function following neonatal HI.

**Methods:** The postnatal day 10 mice were subjected to HI using the Vannucci model. Lipid droplets (LD) were assessed by histology and immunofluorescent staining. Single-nucleus RNA sequencing (snRNA-seq) was performed using brain tissue from HI-injured and sham-operated mice at 72 hours after HI. LD-accumulating microglia (LDAM) were identified by a specific LD marker gene perilipin 2 (*Plin2*). Differential gene expression was analyzed between *Plin2*-positive and *Plin2*-negative microglia after HI. Human HIE brain sections were also examined for LD accumulation. The dynamic changes of PLIN2-expressing microglia and infiltrating monocyte-derived macrophages (MDM) at 24 hours, 72 hours and 7 days after HI were compared using flow cytometry. In addition, mouse BV2 microglia were subjected to oxygen-glucose deprivation (OGD) to study phagocytosis and cytokine expression.

**Results:** Lipid droplets accumulated primarily in microglia after HI in neonatal mice and in human HIE brain. LD were not found in astrocytes or neurons. *Plin2*-expressing LDAM emerged as new microglia clusters after HI. Compared with microglia without LD, LDAM showed a distinct transcriptional profile with upregulation of genes linked to microglial activation, enhanced cholesterol and lipid processing, and a shift towards phagocytic and pro-inflammatory state. Blocking LD biogenesis reduced elevated phagocytosis and IL-1β expression in BV2 cells following OGD.

**Conclusion:** Our study revealed that microglia accumulate lipid droplets as part of their metabolic responses to HI in the neonatal brain. Microglial lipid droplet formation is associated with a pro-inflammatory phenotype at early stage after HI, and increased phagocytosis in vitro. The lipid metabolic changes may regulate microglial function and influence HI outcomes.

## Introduction

Hypoxic-ischemic encephalopathy (HIE), caused by a reduced supply of oxygen and blood flow to the brain, is a leading cause of neonatal mortality. Survivors of HIE often demonstrate persistent motor deficits (such as cerebral palsy, epilepsy) and sensory or cognitive abnormalities [1–3]. Therapeutic hypothermia, while being standard of care for moderate to severe HIE, only provides partial protection and must be initiated within the first 6 hours after birth [4, 5]. Targeting pathways beyond this early time window may provide additional benefit to lower the risk of death and neurodevelopmental impairments.

Inflammation triggers secondary brain damage following initial neuronal death and persists over extended time periods in rodent stroke and HIE models [6]. Hypoxia-ischemia (HI) results in robust microglia activation accompanied by profound metabolic changes. Phagocytic clearance of apoptotic neurons (efferocytosis) is a core function of microglia to reestablish the proper environment for tissue regeneration [7, 8]. We previously reported that at 3 days after HI, microglia, rather than the infiltrated monocyte-derived macrophages (MDM), comprised the major phagocyte population in the neonatal brain [9]. During phagocytosis, microglia experience significant metabolic stress, as they become overloaded with lipids derived from fragmentized membranes, synaptic remnants and myelin debris. Toxic lipid metabolites, including free cholesterol (FC), free fatty acids (FFA), oxidized lipid molecules and other byproducts, are among the major apoptotic cell-derived components following engulfment and lysosomal degradation [10, 11]. It is unclear how microglia process lipid cargo, whether lipid accumulation impacts microglial function, and what the downstream effects are on brain recovery and repair. Recent literature shows that aberrant lipid metabolism, or defective clearance of cholesterol and oxidized lipids is linked to a disease-associated microglial phenotype, and to the development of neurodegenerative diseases as well as impaired remyelination [12–14]. In adult stroke models, lipids, including FC and cholesterol crystals, are observed within microglia in and around the infarct areas [15, 16]. Recently characterized stroke-associated myeloid cells exhibit a lipid-phagocytosing phenotype and are mainly of resident microglial origin [17]. Furthermore, reducing lipid buildup within stroke infarct attenuates chronic inflammation with improved functional outcomes [15]. These results indicate that the ability of microglia to clear excess lipids affects their reactivity and function, directly impacting brain structural and functional recovery. Cells employ protective strategies to prevent lipid buildup by limiting uptake and endogenous synthesis, promoting efflux, and sequestering them within lipid droplets (LD). LD (Fig.1) are unique organelles containing a hydrophobic core of neutral lipids (triglyceride and cholesterol esters) that is enclosed by a phospholipid monolayer and associated proteins (primarily members of the perilipin family) [18–21]. Cholesterol esters (CE) are formed by sterol O-acyltransferase 1 (SOAT1), which catalyzes the esterification of cholesterol with fatty acids for storage in LD [22]. The neutral lipids stored in the LD can be broken down by lipolysis or lipophagy (autophagic degradation of LD) [23, 24]. Lipolysis of triglyceride (TAG) occurs stepwise via three LD surface lipases: ATGL (Adipose Triglyceride Lipase), HSL (Hormone-Sensitive Lipase), and MGL (Monoglyceride Lipase) to produce FAA and glycerol. Free cholesterol released from CE hydrolysis (by lysosomal acid lipase, LAL) during LD lipophagy can be further transported by ATP-binding cassette (ABC) transporter A1 (ABCA1) and G1 (ABCG1) onto ApoE-containing HDL-like particles in the brain [25, 26]. Alternatively, cholesterol can be converted by cholesterol 25-hydroxylaze (CH25H), an enzyme expressed mainly in monocyte lineage cells [27, 28], to 25-hydroxycholesterol (25-HC). As a more mobile oxysterol, 25-HC readily crosses cellular membrane [29–31] and facilitates cholesterol export and clearance (Fig.1). Through dynamic balance of biogenesis and catabolism, LD play key roles in directing the flow of lipid substrates between storage and utilization depending on energy needs. LD biogenesis is common when cells are under metabolic stress; for example, LD-accumulating microglia (LDAM) are prominent in the aging brain and display impaired innate immune responses, leading to worse neurological outcomes after stroke [32]. Aged LDAM are described as proinflammatory and dysfunctional, contributing to age-related forms of neurodegeneration and demyelination disorders [13, 33]. Whether microglia in the developing brain accumulate injury-associated lipids, and how microglial metabolic reprogramming is related to their functionality and contribution to neuroinflammation, remain unknown.

**Fig. 1:**
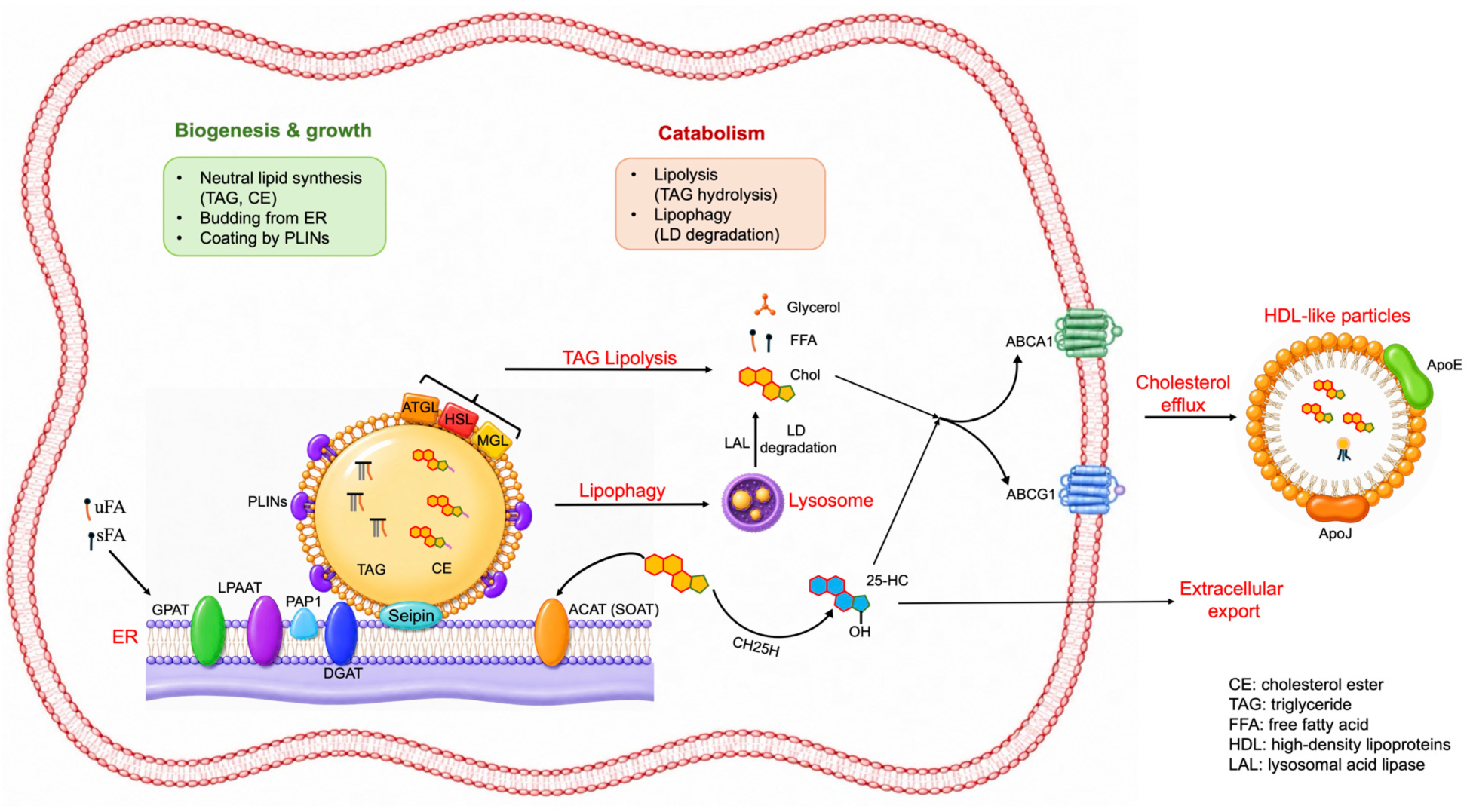
Schematic overview of lipid droplet (LD) biogenesis, turnover and cholesterol handling in microglia. TAG are synthesized in the ER through sequential enzymatic reactions of GPAT, LPAAT, PAP1, and DGAT. CE are generated by SOAT. Seipin facilitates the incorporation of neutral lipids into the growing LD core, which buds from the ER and become coated by perilipins (PLINs). Stored neutral lipids are mobilized through lipolysis mediated by ATGL, HSL, and MGL or degraded via lysosomal lipophagy. Released free fatty acids and cholesterol can be utilized for cellular metabolism, while cholesterol may also undergo ABCA1- and ABCG1-mediated efflux to ApoE-containing HDL-like particles. Alternatively, cholesterol can be converted by CH25H to 25-hydroxycholesterol (25-HC), which readily crosses cellular membranes for export.

In this study, we demonstrate a conspicuous accumulation of LD in microglia at 72 hours (hr) after HI in the neonatal mouse brain. Notably, LDAM are also abundant in human HIE brains. Unlike in the aged brain, LD are generated in neonatal microglia after HI rather than accumulating before the insult, indicating that they are part of the stress response components. In the single-nucleus RNA sequencing (snRNA-seq) experiments, we used perilipin 2 (*Plin2*), a specifically and constitutively expressed LD-associated gene, as a marker gene for LD [34, 35]. We show that *Plin2*-expressing LD-accumulating microglia have a distinct transcriptomic profile characterized by altered cholesterol and lipid homeostasis, and upregulated phagocytic and inflammatory pathways compared to microglia without LD. We suggest a potential link between LD formation and pro-inflammatory cytokine expression after neonatal HI in vivo and in vitro.

## Methods

### Animals

All animal procedures were performed in accordance with the recommendations outlined in the Guide for the Care and Use of Laboratory Animals of the National Institutes of Health, approved by the Institutional Animal Care and Use Committee at University of California, San Francisco (UCSF, #AN202238), and Kunming Medical University (permit# SYXK2015-0002). All surgery protocols were reported in compliance with the Animal Research: Reporting of In Vivo Experiments guidelines. C57BL/6J mice were purchased from the Jackson Laboratory (strain #:000664, Bar Harbor, ME). P2ry12-CreER; Ai14 reporter mice, which express TdTomato upon Cre-dependent recombination, were generously provided by Dr. Thomas Arnold at UCSF [36]. Both sexes were used at postnatal day 10 (P10).

### Human HIE brain sections

Cryosections (16μm) from PFA-fixed OCT-embedded brain tissue (cingulate gyrus) of two HIE patients were obtained from UCSF department of Pathology. Case #1 (UCSF2020-005) was a female, born at 39 gestation weeks with severe HIE and survived 4 weeks. Case #2 (UCSF2021-007) was a female, born at 33 gestation weeks with HIE and survived 7 days.

### Neonatal brain Hypoxia-ischemia (HI)

Unilateral HI was induced using the modified Vannucci model in P10 pups as we published [9, 37–39]. Through a vertical midline neck incision under isoflurane anesthesia (2-3% isoflurane in balanced oxygen), the left common carotid artery (CCA) was permanently occluded using electrical coagulation. The animals were allowed to recover for 1hr with their dam and then exposed to 45 min of global hypoxia (8% oxygen balanced with nitrogen) in a humidified chamber at 36.5 °C. Sham-operated control animals received isoflurane anesthesia and exposure of CCA without coagulation and hypoxia. In HI animals, the left ipsilateral hemisphere experiences both hypoxia and ischemia, whereas the right contralateral side experiences hypoxia alone. Our procedure produces moderate to several injury in the ipsilateral hemisphere, primarily in the cortex, hippocampus, striatum and thalamus as shown in Fig. 3 and Fig. 4.

**Fig. 2:**
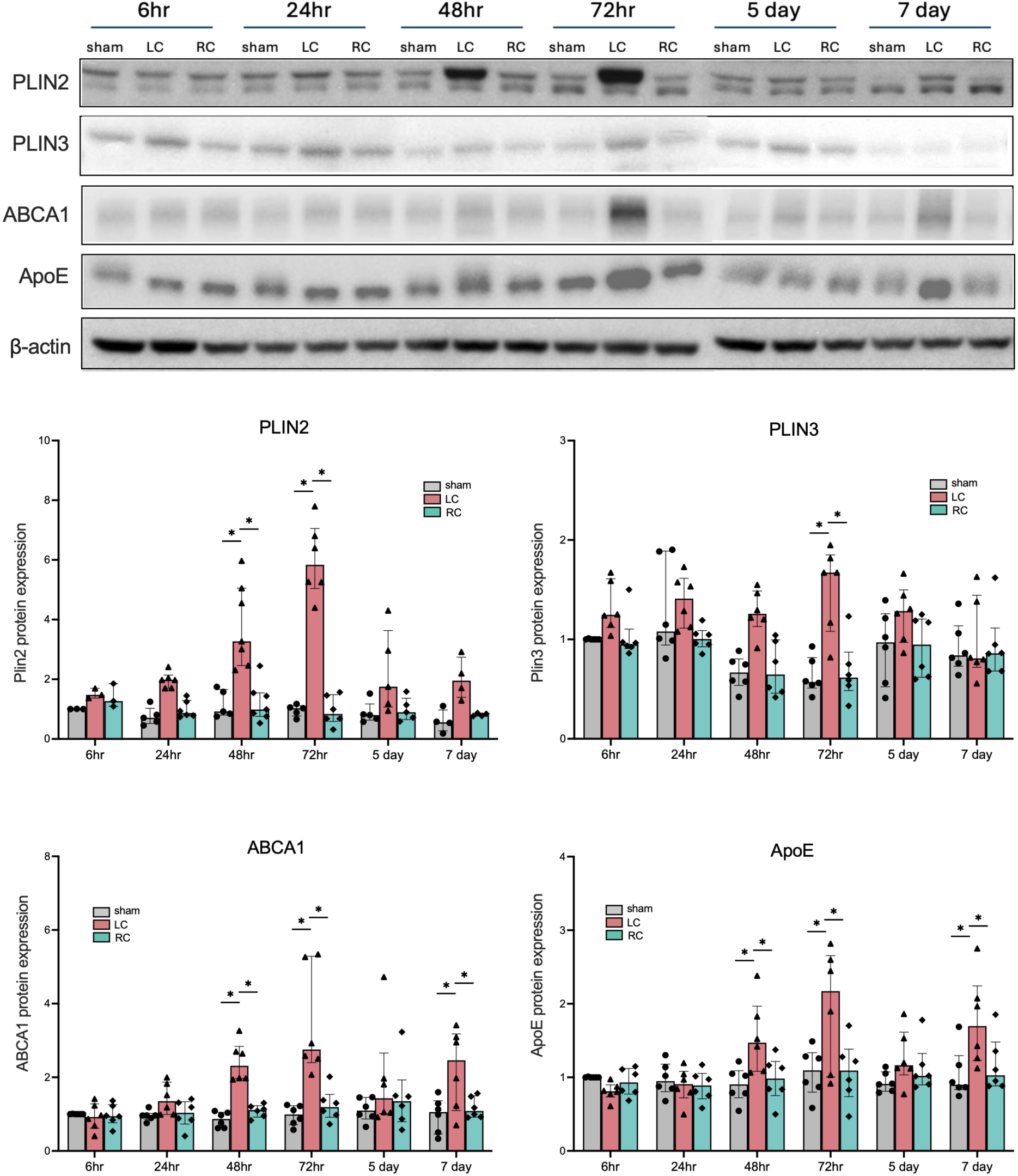
Upregulation of lipid droplets-associated protein expression after HI in neonatal mouse brain. The representative Western blotting images of PLIN2, PLIN3, ABCA1, ApoE and β-actin from 6hr to 7 days after HI are shown at the top and the quantification (normalized to β-actin and then to the value of sham 6hr) are shown at the bottom. n=3-5 for sham, n=4-6 for LC: left cortices, the ipsilateral side; and RC: right cortices, the contralateral side. *: p<0.05

**Fig. 3:**
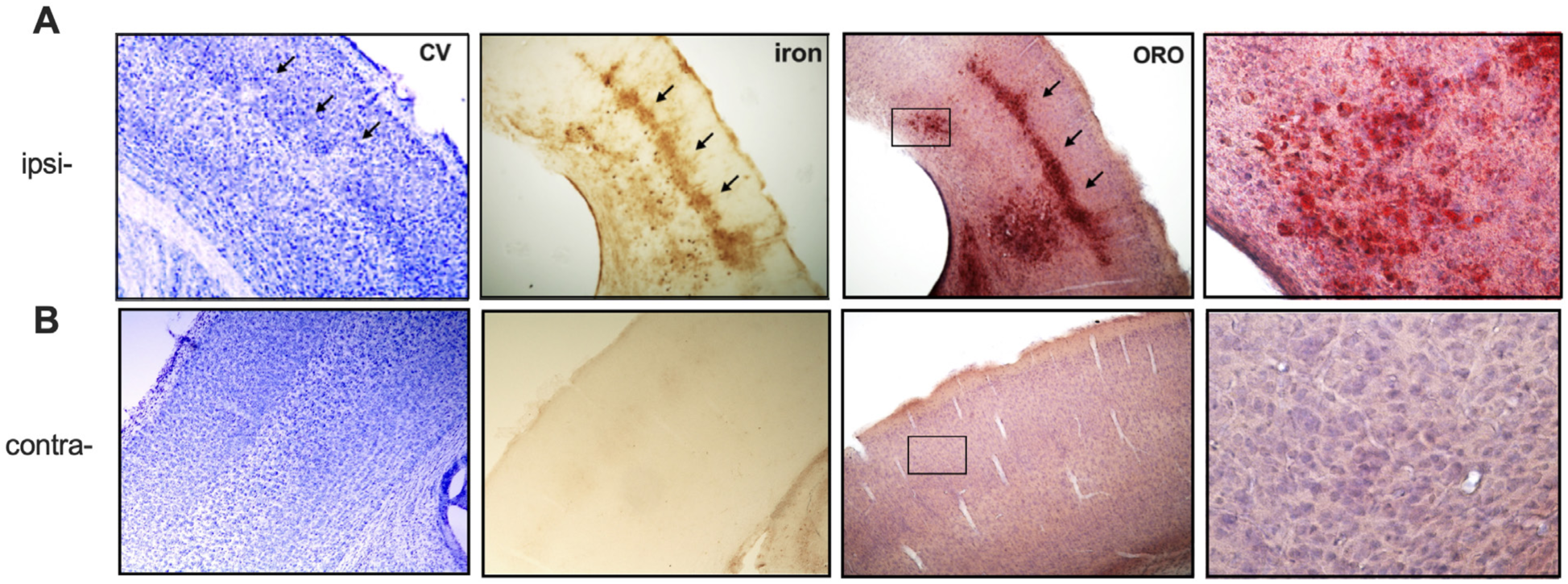
Accumulation of Lipid droplets after HI in neonatal mouse brain. The representative images of cresyl violet (CV), iron and ORO staining are shown in the ipsilateral (ipsi-) cortex (A) and the contralateral (contra-) cortex (B). A). The pattern of red ORO staining (arrows) corresponded to that of iron deposit (arrows) and areas of cell loss (arrows) shown with CV staining. B). Cell loss, iron deposit and ORO staining were not observed in the contralateral cortex. The sections were counterstained with hematoxylin in the ORO images.

**Fig. 4:**
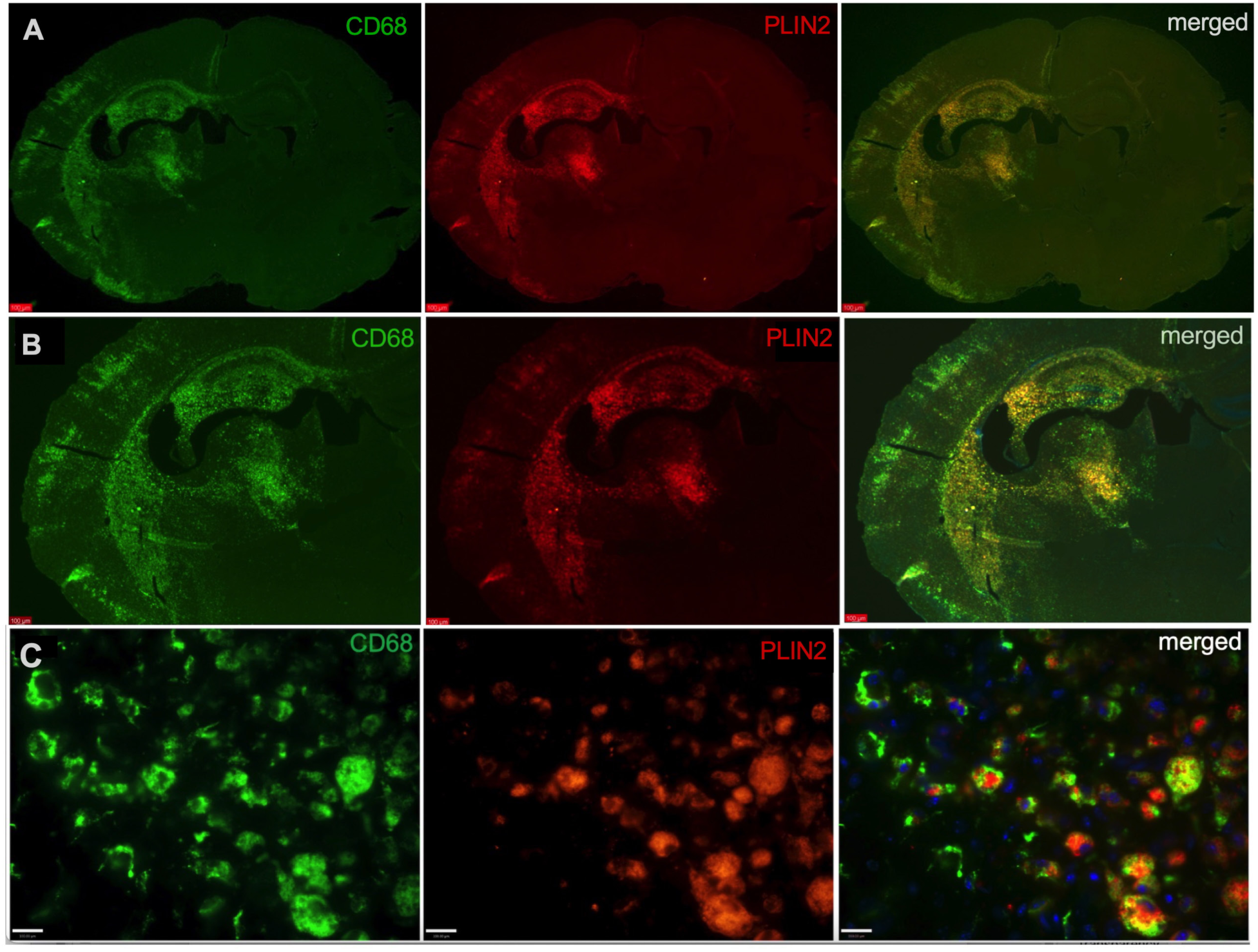
Lipid droplets build up in activated CD68+ microglia/MDM at 72hr after HI. A) and B): double immunofluorescent staining shows colocalization of CD68 (green) and PLIN2 (red) in the ipsilateral hemisphere at 72hr after HI in the typical injured brain regions (cortex, hippocampus, striatum and thalamus). CD68 and PLIN2 were not expressed in the contralateral hemisphere. C): high magnification images (63x) show foamy CD68+/PLIN2+ double positive cells (n=3).

### Histology staining

Animals were transcardially perfused with cold PBS followed by 4% paraformaldehyde (PFA). Brains were dissected, post-fixed for 24hr in 4% PFA at 4°C, and cryoprotected in 30% sucrose. 50-μm-thick serial vibratome sections were stained with cresyl violet (CV) and iron as we previously published [37, 39, 40].

Oil Red O (ORO) stains neutral lipids, most commonly cholesterol ester and TAG, which form the hydrophobic core of LD. Therefore, ORO staining is widely used to visualize LD in tissue sections and cultured cells. ORO (Cat# O-0625, Sigma-Aldrich, Inc., St. Louis, MO) stock solution (0.35g ORO in 100ml of isopropanol) was filtered and stored at room temperature (RT). The working solution was prepared freshly by mixing 3 parts of the stock solution with 2 parts of ddH_2_O. Brain sections were incubated with ORO working solution at RT for 10 minutes (in the dark) and washed once with 60% isopropanol for 30 seconds followed by ddH_2_O wash 2-3 times. The sections were counterstained with hematoxylin for 1 minute and coverslipped with warmed glycerol gelation.

### Western Blotting

Cortical protein from the sham and HI-injured animals (both contralateral and ipsilateral side) was extracted with RIPA lysis buffer (Cat# R0278, Sigma) containing Halt^TM^ proteinase and phosphatase inhibitors (Cat# 78442, Pierce Biotechnology, Rockford, IL). Western blotting was performed using the standard protocol, with the following primary antibodies: PLIN2 (1: 500, Cat# NB110-40877, Novus Biologicals, LLC, Centennial, CO); PLIN3 (1:1000, Cat# sc-390968, Santa Cruz Biotechnology, Inc, Dallas, TX); ABCA1(1:1000, Cat# 96292S, Cell Signaling Technology, Inc., Danvers, MA), and ApoE (1:1000, Cat# 49285S, Cell Signaling Technology). The membranes were stripped and reprobed with mouse β-actin antibody (1:6000, sc-47778, Santa Cruz Biotechnology, Inc.). Appropriate HRP-conjugated secondary antibodies were used and protein signal was visualized with enhanced chemiluminescence with ChemiDoc Imaging System (Bio-Rad Laboratories, Hercules, CA). Image J software was used to measure the mean optical density (OD) and the area of protein bands. OD values of each protein were normalized to β-actin and presented as the fold changes relative to sham controls at 6hr time point.

### Immunofluorescent staining of mouse HI and human HIE brain sections

Mouse HI brain cryosections (16μm) were washed in PBS followed by blocking with PBS containing 10% donkey serum and 0.25% Triton X100 at RT for 1 hr. Sections were incubated at 4°C overnight with rabbit anti-PLIN2 antibody (1:200, Cat# 15294-1-AP, Proteintech Group, Inc, Rosemont, IL) paired with another antibody that is specific to neurons (mouse anti-NeuN, 1:400, Cat# MAB377, EMD Millipore, St. Louis, MO) or astrocytes (chicken anti-GFAP, 1:1000, Cat# PA1-10004, Invitrogen, Carlsbad, CA) or activated microglia/macrophages (rat anti-CD68, 1:300, Cat# MCA1957, Bio-Rad). The antibodies were diluted in PBS containing 5% donkey serum and 0.25% Triton X100. After washing, the secondary antibodies (corresponding cross-adsorbed Alexa Fluor antibodies; Invitrogen) were applied for 1 hr at RT. Following washing with PBS, the slides were coverslipped with DAPI Fluoromount-G mounting medium (Cat# 0100-20, SouthernBiotech, Birmingham, AL).

For the P2ry12-CreER^/+^; Ai14^/+^ heterozygotes, tamoxifen (8mg/ml in corn oil, Cat# T5648, Sigma) was injected intraperitoneally at 50mg/kg at P6 and P7 to induce recombination and expression of TdTomato (TdT). The mice were subjected to HI at P10. At 72hr after HI, the brains were sectioned and stained with the same rabbit anti-PLIN2 antibody. Staining for human brain cryosections was performed as described for mouse sections using the same rabbit anti-PLIN2 antibody combined with goat anti-Iba1 antibody (1:300, Cat# NB100-1028, Novus) or mouse anti-NeuN or chicken anti-GFAP antibody (same as above).

### BODIPY^TM^ 493/503 staining of mouse HI and human HIE brain sections

BODIPY™ 493/503 is a fluorescent probe that specifically labels neutral lipids and is commonly used to detect LD [41]. BODIPY^TM^ 493/503 (Cat# D3922, Thermo Fisher Scientific, Waltham, MA) stock solution was made in ethanol at 0.2 mg/ml. Brain cryosections from the P2ry12-CreER^/+^; Ai14^/+^ mice, and HIE human brain were washed 3 times with PBS following the secondary antibody staining, and incubated with BODIPY (1:400 dilution from the stock) for 15 mins at RT. After washing, the slides were coverslipped with DAPI Fluoromount-G mounting medium.

Images were captured using an Olympus FLUOVIEW FV3000 confocal microscope and processed using Fiji/ImageJ (NIH) software.

### Flow cytometry analysis of mouse HI brain samples

Following transcardial perfusion with ice-cold 0.9% saline, cortical single cell suspensions were prepared from C57BL/6J mice at 24hr, 72hr and 7 days after HI or sham surgery, using Neural Tissue Dissociation Kit (Cat# 130-092-628, Miltenyi Biotec, Inc. San Jose, CA) as we published [9]. Myelin debris was removed with Myelin Removal Beads (Cat# 130-096-733, Miltenyi Biotec, Inc.). Cells were first stained with Fixable Viability Dye eFluor™ 780 (1:1000; Cat# 65-0865-14, eBioscience) to exclude dead cells and then blocked with anti-CD16/32 antibody (1:100, Biolegend, San Diego, CA). The cells were incubated with a surface antibody mixture of APC anti-mouse CD45 (1:100, Cat# 157606, Biolegend) and BV785 anti-mouse CD11b (1:100, Cat# 101243, Biolegend) antibodies at RT for 30 min in the dark. After washing, cells were fixed and permeabilized with BD Fixation/Permeabilization Kit (Cat# 554722, BD Biosciences, San Jose, CA), followed by intracellular staining with rabbit anti-PLIN2 (1:200, Cat# PA1-16972, Invitrogen), PE anti-IL-6 (1:100, Cat# 504503, Biolegend), and PE-cyanine7 anti-IL-1β (1:100, Cat# 25-7114-82, eBioscience) at 4°C for 30 min. For PLIN2 detection, cells were additionally incubated with donkey anti-rabbit Alexa Fluor 647 (1:200, Cat# ab150075, Abcam) at 4°C for 30 min in the dark. After washing, cells were resuspended in 300 µL of DPBS with 0.5% FBS and analyzed on a 38-color spectral flow cytometer (Cytek Aurora, Cytek Biosciences Co., Fremont, CA). Compensation beads (BD Biosciences) were used to establish the spectral unmixing matrix. Fluorescence-minus-one (FMO) controls and isotype-matched control antibodies were included for all intracellular staining to ensure accurate gating. At least 1.5 × 10⁵ viable cells were acquired per sample, and data were analyzed using FlowJo software (version 10.8.1; Tree Star Inc. Ashland, OR). Microglia were defined as CD45^int^CD11b^+^ cells, while infiltrating monocytes-derived macrophages (MDM) were defined as CD45^high^CD11b^+^ cells. The same gating strategy was applied to all groups. All experiments were independently repeated at least three times.

### Mouse brain single-nucleus RNA sequencing (snRNA-seq)

At 72hr after HI, the ipsilateral cortices from three animals per group (HI- or sham-operated) were used for the snRNA-seq experiments.

(1). Single-cell samples were prepared as described in the flow cytometry experiments. The cells of three animals per group were pooled for further analysis.
(2). Nucleus acquisition and quality control: cell nuclei were isolated using the Chromium Nuclei Isolation Kit with RNase Inhibitor (Cat# PN-1000494, 10x Genomics, Pleasanton, CA). After filtering through a 70μm cell strainer, washing and centrifugation, the cell pellets were resuspended in 100 μl ice-cold Nucleus Storage Buffer (NSB: 1× PBS, 1% BSA and 0.2 U/µL RNase Inhibitor, Takara). After quality control, samples with >70% nuclear integrity, <5% fragmentation, and no significant clumping were used, and adjusted to a concentration of 1,000 nuclei/μl for subsequent library preparation.
(3). 10x Genomics library construction and sequencing: The purified cell nuclei were separately loaded on the Chromium Single Cell Controller (10x Genomics) using the Chromium Next GEM Single Cell 3’ Reagent Kits (v3.1, Cat# PN-1000268) according to the manufacture’s protocol. Single-nuclei suspensions were encapsulated into droplets, with the RNA from each nucleus reverse transcribed using a unique oligonucleotide barcode. The gel beads, single nuclei suspension and the master mix were loaded onto Chromium Next GEM Chip G. Reverse transcription was performed at 53°C for 45 min followed by 12 cycles of cDNA amplification. Fragmented products (∼400 bp) underwent 14 cycles of index PCR to generate the final library. Library concentration was quantified using the Qubit 4 dsDNA HS Assay (Thermo Fisher Scientific Inc.), and fragment distribution was confirmed by the 4200 TapeStation system (Agilent Technologies, Santa Clara, CA). The libraries were pooled equimolarly to 2 nM. Sequencing was performed on a NovaSeq^TM^ 6000 using SP flow cells in 28-8-0-91 mode, with a sequencing depth of ≥ 50,000 reads per nucleus.

### snRNA-seq data analysis

Preprocessing and quality control of the snRNA-seq data were performed using the Cell Ranger toolkit (v6.1.1, 10x Genomics) [42]. The ‘mkfastq’ pipeline was used for demultiplexing, and the ‘cell ranger count’ pipeline was used to map the obtained FASTQ file to the mouse reference genome mm10, generating a gene-barcode matrix.

Downstream analysis of the matrix was performed using the Seurat (v5.4.0) R package [43]. Cells were filtered based on number of expressed features and mitochondrial genes proportion, retaining only cells with 300-7000 RNA features and < 10% mitochondrial gene composition. Gene expression values were log-normalized and scaled to 10,000 transcripts per nucleus. Top 3000 highly variable genes (HVGs) were selected using the variance stabilizing transformation (vst) method. 50 principal components (PCs) were used for principal component analysis (PCA). Dimension reduction was later performed using uniform manifold approximation and projection (UMAP) and t-distributed stochastic neighbour embedding (t-SNE).

Cell populations were identified based on known markers genes. Given the focus of this study is microglia, microglia population was identified based on the expression of *P2ry12* [44]*, Cx3cr1* [45]*, Itgam* and *C1qa,* and subsetted from other cell types (Supplementary Fig.1). The subset was re-normalized and scaled, and the top 2000 HVGs were selected. 44 PCs, cumulatively accounted for 90% of the total variance, were used for PCA. Dimension reduction was performed using UMAP and t-SNE. At resolution 1.0, a reasonable number of clear clusters were identified. Seurat’s FindAllMarkers function was used to identify cluster-specific marker genes (log2 fold-change ≥ 1, minimum expression percentage: 0.3). Based on expression profiles and published literature, clusters were merged into six clusters (Supplementary Fig. 2). Visualization was performed using R packages ggplot2 (v4.0.1) and Seurat (v5.4.0).

### Differentially Expressed Genes and Functional Analysis

Differential expression analysis between experimental groups were performed using the non-parametric Wilcoxon rank sum test with Benjamini-Hochberg false discovery rate (BHFDR adjusted p-value (Padj) < 0.05). Genes were defined as differentially expressed (DEGs) if Padj < 0.05 and |log_2_ fold change (FC)| > 1). Pathway analyses were performed using the R package clusterProfiler (v4.8)[46], with significance set at Padj < 0.05. Heatmaps and volcano plots were generated to visualize the results.

Gene Ontology (GO) enrichment analysis was performed using gene set sizes of 10-500 genes, and a background of genome-wide protein-coding genes from the mm10 reference genome, including Biological Process (BP), Molecular Function (MF), and Cellular Component (CC). KEGG and Hallmark gene set enrichment analyses were also conducted. Results were visualized using clusterProfiler (v4.8) and ggplot2 (v4.0.1).

Sequencing data have been deposited in the Open Archive for Miscellaneous Data (https://ngdc.cncb.ac.cn/omix/: accession no: OMIX014659).

### BV2 Cell Culture and Oxygen-Glucose Deprivation (OGD)

BV2 murine microglial cells were obtained from the Cell Bank of the Chinese Academy of Sciences. Cells were plated at 1 × 10^6^ cells per 25 cm^2^ flask with high-glucose DMEM with 8% FBS and 1% penicillin/streptomycin. All experiments were performed using BV2 cells at passages 3 to 10.

For OGD experiments, cells were trypsinized and seeded at 2 × 10^4^ cells per 14-mm^2^ coverslip, or 1 × 10^5^ cells per well in 12-well plates. After 12 hours, cells were washed with PBS and the culture medium was replaced with glucose-free DMEM (Cat# 90113, Solarbio, China). OGD was induced by placing cells in a humidified Modular Incubator Chamber (Cat# MIC-101, Billups-Rothenburg Inc. San Diego, CA) flushed with 95% N_2_ / 5% CO_2_ for 3 min at 40 L/min and incubating at 37°C for 2 hours. OGD was terminated by replacing the medium with DMEM containing 8% FBS, and the cells were cultured for an additional 24 hours. Control cells were maintained in normal DMEM under normoxic condition.

DGAT1 inhibitor T863 (Cat# HY-32219, MedChemiExpress LCC, Monmouth Junction, NJ), and DGAT2 inhibitor PF-06424439 (Cat# HY-108341A, MCE) were dissolved in DMSO and used at 30 µM and 40 µM, respectively. BV2 cells were pretreated with the inhibitors for 12 hours before OGD, and the drug was maintained during OGD and the 24-hr reoxygenation period. Control cells received an equivalent concentration of DMSO (≤ 0.1%) as vehicle control.

### Phagocytosis Assay using FITC-Dextran 40 in BV2 microglial cells

After OGD, BV2 cells were incubated with FITC-labeled 40-kDa dextran (1mg/ml, Cat# FD40, TdB Labs AB, Sweden) at 37°C for 30 min. Cells were washed twice with PBS to remove unbound FITC-dextran 40, coverslipped with Fluoroshield mounting medium with DAPI (Cat# F6057, Sigma) and imaged under a fluorescence microscope (Zeiss).

Flow cytometry was also performed to quantitatively assess FITC-Dextran 40 uptake by BV2 cells. After FITC-dextran 40 incubation and subsequent washing, BV2 cells were trypsinized, washed, and stained with Fixable Viability Dye eFluor™ 780 (1:1000; Cat# 65-0865-14, eBioscience) to exclude dead cells. After washing, cells were analyzed using a flow cytometer (Cytek Aurora). At least 1 × 10⁴ viable cells were acquired per sample. Data were analyzed with FlowJo software.

### BODIPY staining and flow cytometry analysis for BV2 cells

To evaluate DGAT inhibitor efficacy, lipid droplet formation was assessed as a readout. To visualize LD, BV2 cells were washed twice with ice-cold PBS and incubated with BODIPY 493/503 (1:1000; Cat# C2053S, Beyotime, China) at 37°C for 20 min in the dark. After washing with PBS, cells were coversliped with Fluoroshield mounting medium with DAPI and imaged using a confocal laser scanning microscope (Zeiss, LSM 800, Germany).

Flow cytometry was performed to evaluate cytokine expression in LD-containing BV2 cells. BODIPY stained cells were trypsinized, washed with ice-cold PBS, and incubated with Fixable Viability Dye eFluor™ 780 to exclude dead cells. Cells were then stained with antibody against PE-cyanine7 IL-1β as described above in the flow cytometry analysis with mouse brain samples. Cells were analyzed on Cytek Aurora flow cytometer. Gating and data analysis were performed using FlowJo software.

### Immunofluorescence staining of BV2 microglial cells

Following OGD, for double staining of PLIN2 and FITC-dextran 40, cells were first incubated with FITC-dextran 40, washed and then fixed with 2% paraformaldehyde for 15 min. Cells were permeabilized with 0.3% Triton X-100 for 15 min and blocked with 5% donkey serum at RT for 1 hour. Cells were incubated with rabbit anti-PLIN2 (1:200; Cat# PA1-16972, Invitrogen) overnight at 4°C. After washing with PBS, cells were incubated with donkey anti-rabbit Alexa Fluor 647 (1:200; Cat# ab150075, Abcam,) at RT for 1 hour, washed and coverslipped. For double staining of BODIPY and IL-1β, cells were first stained with BODIPY 493/503, fixed followed by staining with anti-IL-1β primary antibody (1:100; Cat# GTX100793, Gentex, Zeeland, MI) using the same protocol as above.

### Statistical Analysis

GraphPad Prism 9.2 was used for graph generation and statistical analyses. For Western blotting and brain 25-HC concentrations, comparisons between two groups (values of sham vs. ipsi- or contralateral cortex, ipsilateral vs. contralateral cortex at each individual time point) were performed using nonparametric Mann-Whitney test. Data are presented as median with interquartile range using interleaved scatter plots with bars. For flow cytometry analyses, two-way ANOVA with Tukey’s multiple comparisons test was used. The specific statistic method was included in the individual figure legend. For all results, differences were considered statistically significant at p< 0.05. Statistical methods for snRNA-seq and DEG analyses were described above.

## Results

### Upregulation of lipid droplet-associated protein after HI in neonatal mouse brain

While PLIN2 and PLIN3 are both LD-coating proteins, PLIN2 is the best-characterized, more specific and widely used marker for LD [47, 48]. To determine the time course of PLIN2 and PLIN3 protein expression, as an indication of LD accumulation, we performed Western blotting using cortical lysates from 6hr to 7 days after HI (Fig. 2). The PLIN2 protein levels in the left cortices (LC, the ipsilateral side) increased over time and peaked at 72hr post-HI, after which its expression gradually declined to levels comparable to those of the sham animals. PLIN2 levels remained unchanged in the right cortices (RC, the contralateral side). Similarly, PLIN3 expression was significantly elevated at 72hr in LC compared to the sham and RC values. The expression of ABCA1 and ApoE, two proteins responsible for cholesterol efflux, was increased at 48hr, 72hr and remained higher at 7 days after HI (Fig. 2).

### Accumulation of lipid droplets following HI in neonatal mouse brain

Serial brain vibratome sections were stained with ORO to visualize LD, cresyl violet for overall brain structure, and iron, which is normally deposited in the injured brain regions as we reported previously [37, 39, 40]. Fig. 3 shows that at 72hr after HI, the red ORO-stained regions (arrows in Fig. 3A) corresponded to areas of iron deposition and the regions of columnar cell loss in the cortex (Fig. 3A, arrows). The staining was present only in the ipsilateral hemisphere (Fig. 3A), but not in the contralateral side (Fig. 3B) of HI animals. It has been reported that iron accumulation is a prominent feature of activated microglia in response to inflammation [49–52].

### Lipid droplets are built up predominately in activated microglia after neonatal HI

To determine the cellular distribution of LD, we performed double immunostaining with antibodies against PLIN2, paired with anti-NeuN or anti-GFAP or anti-CD68 (for activated microglia and infiltrating MDM) antibodies. Strikingly, immunoreactivity of CD68 and PLIN2 overlapped in the injured regions of the ipsilateral hemisphere (Fig. 4A, 4B). The abundant double PLIN2+/CD68+ cells suggest LD accumulation in activated microglia (and/or MDM). Under higher magnification (63x), these cells were large, amoeboid and foamy, resembling foam cells in arteriosclerosis (Fig. 4C).

As CD68 is a shared marker for other monocyte/macrophage lineage cells, we used P2ry12-CreER; Ai14 reporter mice, in which TdT expression is driven by *P2ry12* promoter that is specific for parenchymal microglia [36]. Fig. 5A shows that PLIN2 was expressed in TdT-positive microglia in the HI-injured ipsilateral cortex. Microglia in the contralateral side possessed a highly ramified morphology without PLIN2 expression. Consistently, BODIPY staining was localized predominantly in TdT-positive microglia (Fig. 5B) with very few BODIPY^+^/TdT^-^ cells.

**Fig. 5:**
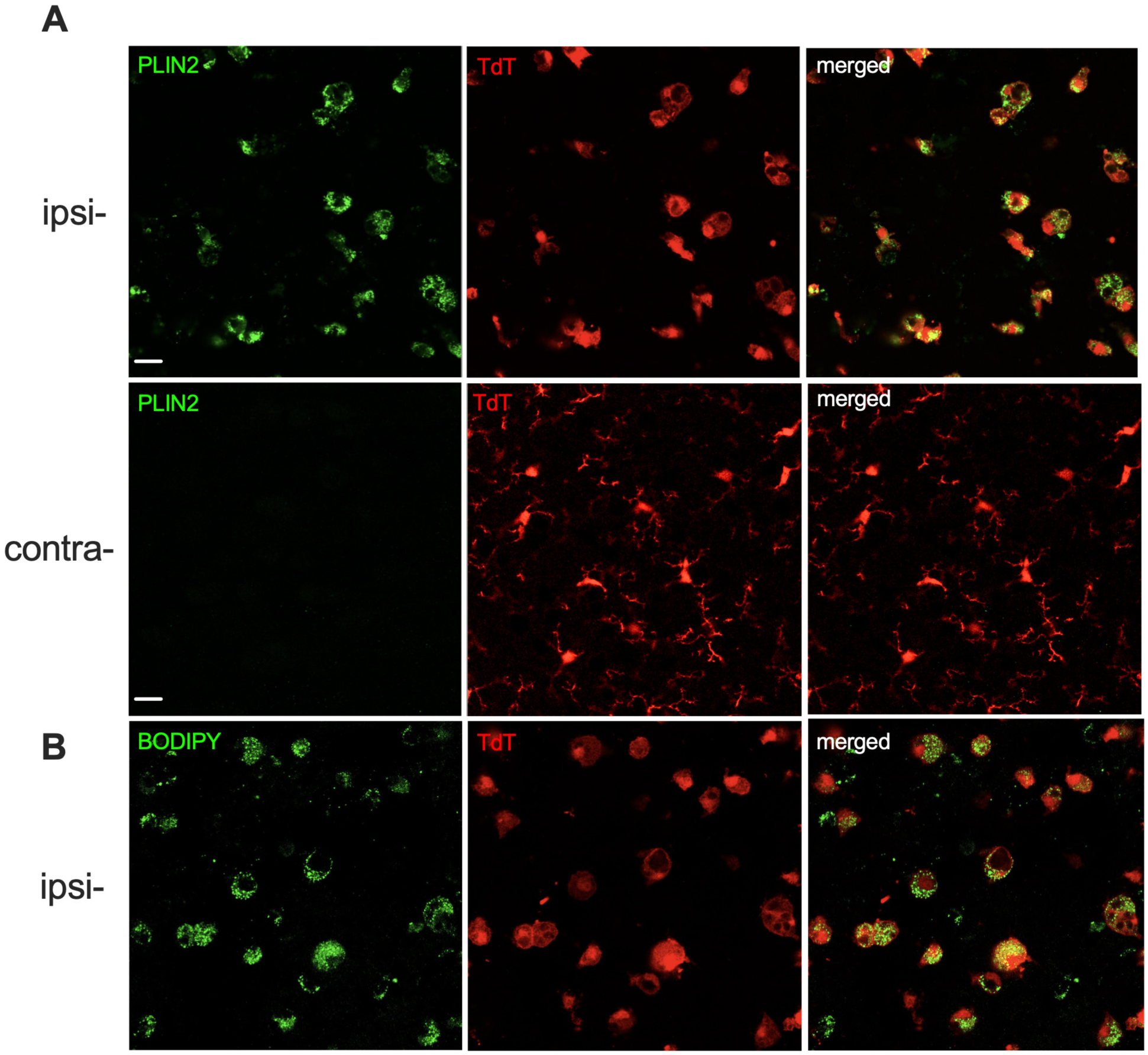
Accumulation of lipid droplets in microglia in P2ry12CreER^/+^;Ai14^/+^ reporter mice after HI. A). PLIN2 (green) was expressed in TdTomato (TdT)-positive microglia in the ipsilateral (ipsi-), but not in the contralateral (contra-) cortex at 72hr after HI. Amoeboid microglia were prominent in the ipsilateral cortex, while they were highly ramified in the contralateral cortex (n=3). B). BODIPY 493/503 (green)-positive lipid droplets were also localized predominantly in TdT+ microglia.

To investigate the temporal dynamics of lipid droplet accumulation in microglia, and whether MDM also accumulate lipid droplets, especially at early time points following HI, we performed flow cytometry analysis in which microglia and MDM were distinguished based on CD11b and CD45 expression. As shown in Fig. 6B, the percentage of PLIN2+ microglia (CD45^int^CD11b^+^) among total number of microglia was significantly elevated in all HI groups compared to their respective sham groups. There was a higher proportion of microglia expressing PLIN2 at 3 days than at 1 and 7 days after HI. LD formation was not restricted to microglia. We calculated the percentage of PLIN2+ microglia (CD45^int^CD11b^+^) and PLIN2+ MDM (CD45^high^CD11b^+^) out of the total number of myeloid cells (CD45^+^CD11b^+^). PLIN2-positive cells were consistently more abundant among microglia (∼ 2 to 2.5 folds higher) than MDM (Fig. 6C). The percentage of PLIN2+ microglia peaked at 3 days after HI and remained significantly higher than PLIN2+ MDM at day 7 (Fig. 6C). The PLIN2+ MDM declined over time in the first week after HI. These results suggest that both microglia and MDM accumulate lipid droplet after injury, but microglia are the main contributor to LD-accumulating population following neonatal HI. LD were not present in NeuN-positive neurons or GFAP-positive astrocytes (Supplementary Fig. 1). Fig. S1A and S1B show that PLIN2 was expressed in the ipsilateral, but not in the contralateral cortex at 72hr after HI. Neurons and astrocytes demonstrated dramatic morphological changes after HI. In the ipsilateral cortex, neurons were significantly shrunken suggesting cell death, while astrocytes had pronounced hypertrophy of the cell body and processes indicating a reactive status as compared to the astrocytes in the contralateral side. No colocalization of PLIN2 with NeuN or GFAP was observed.

**Fig. 6:**
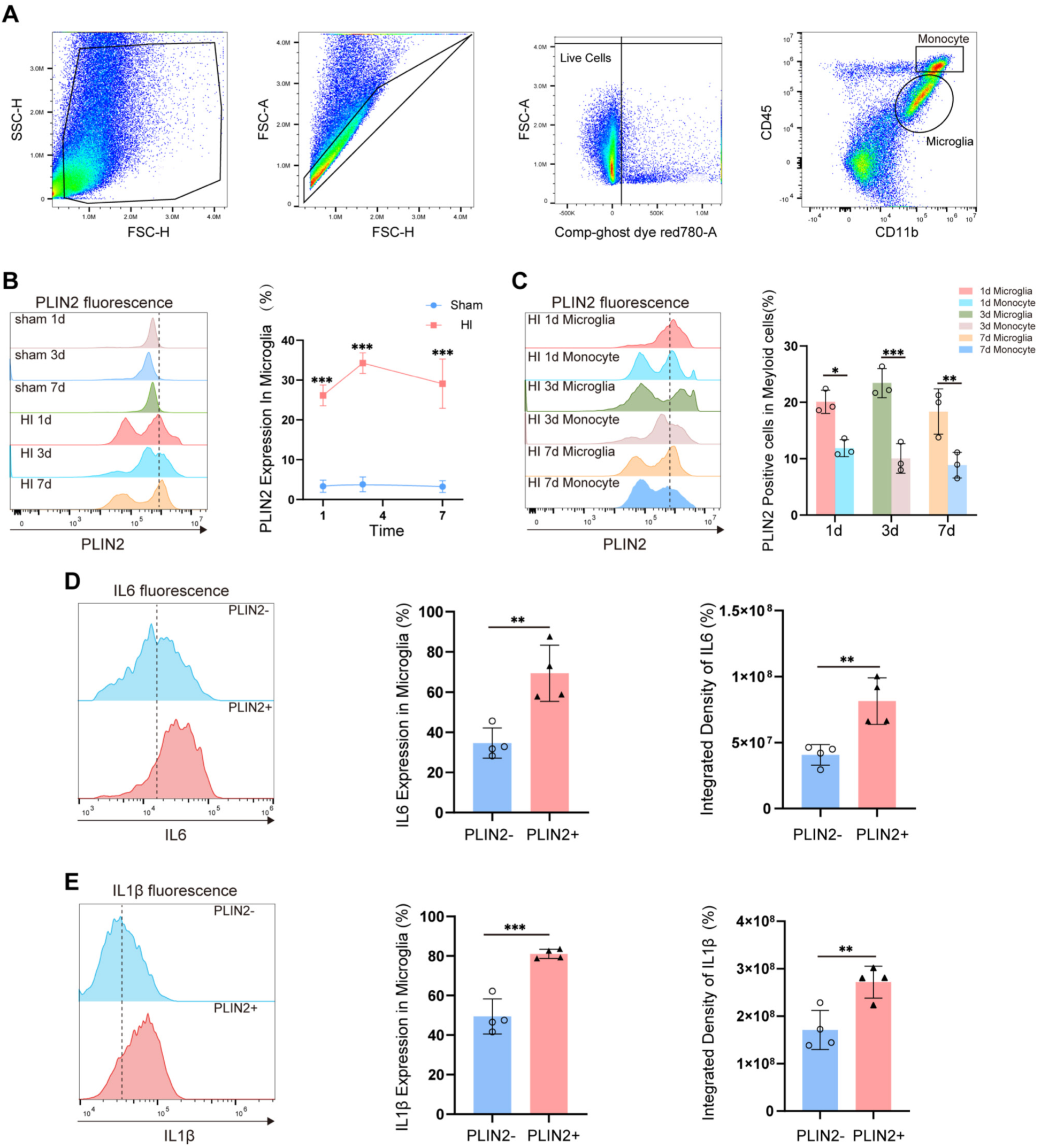
Temporal dynamics of lipid droplet accumulation in microglia and infiltrating monocyte-derived macrophages (MDM) within one week after HI. A). Gating strategy for the identification of microglia and MDM by flow cytometry. Cells were first gated on the FSC-H/SSC-H scatter plot to exclude debris and cell aggregates, followed by singlet selection based on FSC-A/FSC-H. Viable cells were then selected as Ghost Dye Red 780-negative. Microglia and MDM were distinguished based on CD11b and CD45 expression, with CD45^int^CD11b^+^ cells defined as microglia and CD45^high^CD11b^+^ cells as MDM. B). Temporal dynamics of the percentage of PLIN2^+^ microglia at different time points after HI. Left: representative histogram overlays showing PLIN2 fluorescence intensity curves in microglia at each time point. Right: percentage of PLIN2-positive microglia among total microglia in the ipsilateral cortex at 1, 3 and 7 days in the HI and sham groups. (n = 3 per group per time point); two-way ANOVA (time × treatment) with Tukey’s multiple comparisons test; ***p < 0.001 vs. corresponding sham group). C). Histogram and quantification of the percentage of PLIN2^+^ microglia and PLIN2^+^ MDM among total myeloid cells at different time points after HI. (n = 3 per group per time point; unpaired Student’s t-test). D). IL-6 expression in PLIN2^+^ and PLIN2^-^microglia at 3 days after HI. Left: IL-6 fluorescence intensity in PLIN2^+^ (red) versus PLIN2^-^ (blue) microglia. Middle: percentage of IL-6^+^ cells within the two subsets. Right: integrated IL-6 fluorescence (mean fluorescence intensity (MFI) × cell number within each subset) between PLIN2^+^ and PLIN2^-^ microglia. (n = 4 per group, paired Student’s t-test). E). IL-1β expression in PLIN2^+^ and PLIN2^-^ microglia at 3 days after HI. Left: IL-1β fluorescence intensity in PLIN2^+^ (red) versus PLIN2^-^ (blue) microglia. Middle: percentage of IL-1β^+^ cells within the two subsets. Right: integrated IL-1β fluorescence between PLIN2^+^ and PLIN2^-^ microglia. (n = 4 per group). All data are presented as mean ± SD; *p < 0.05, **p < 0.01, ***p < 0.001

### Lipid droplets are present in microglia of human HIE brain

In two HIE cases, we observed abundant Iba1-positive microglia co-expressing PLIN2 or BODIPY in the injured areas of cingulate gyrus (Fig. 7) indicating LD accumulation. These microglia were noticeably ameboid or hypertrophic, suggesting a reactive state. PLIN2 was not expressed in the non-injured areas (Fig. 7D), where microglia were highly ramified. LD were not observed in astrocytes or neurons in human HIE brains (Supplementary Fig. 2). These observations were consistent with the findings in our mouse neonatal HI model.

**Fig. 7:**
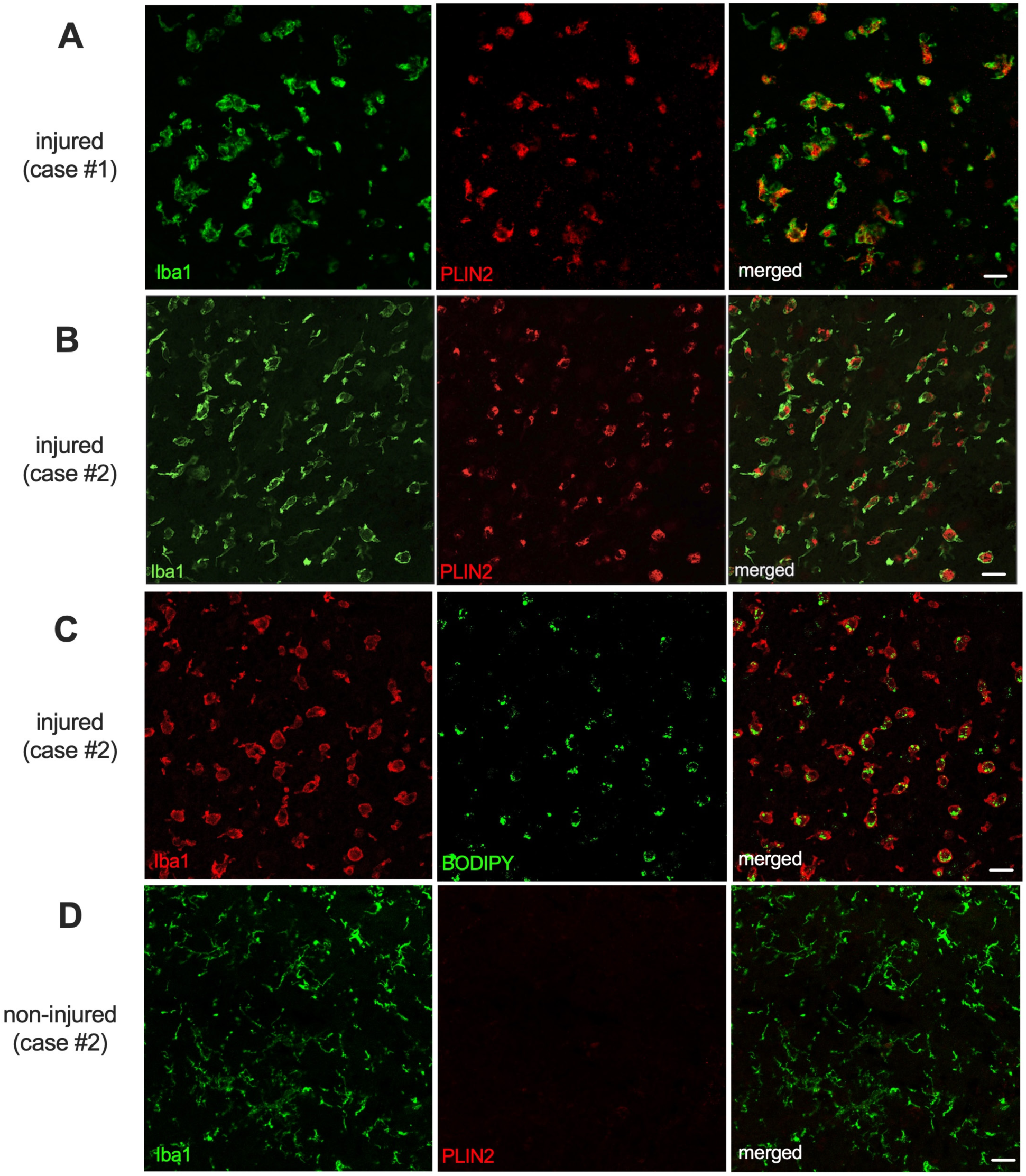
LD-accumulating microglia in human HIE brain. A) and B). PLIN2 (red)-expressing lipid droplets were localized in Iba1-positive (green) microglia in the injured areas of cingulate cortex in two human HIE cases (case#1: female, born at 39 gestation week, diagnosed with severe HIE and survived for 4 weeks; case#2: female, born at 33 gestation week, diagnosed with HIE and survived for 7 days). C). BODIPY (green)-positive LD were localized in Iba1-positive (red) microglia. D). PLIN2 (red) was not expressed in the non-injured areas where microglia were highly ramified.

### New microglia subsets emerge after neonatal HI

To study microglia diversity and responses to HI, we performed snRNA-seq using sham and HI-injured ipsilateral cortices at 72hr post-HI. Microglia clusters were extracted based on canonical homeostatic marker genes (*P2ry12, Cx3Cr1, Itgam* and *C1qa*, Supplementary Fig. 1). We identified new microglial subclusters that emerged after HI, primarily cluster 3 and 4 in Fig. 8A, indicating that these cells acquired distinct gene expression profiles in response to HI compared to microglia in sham animals, in which the cluster 1 microglia predominated.

**Fig. 8:**
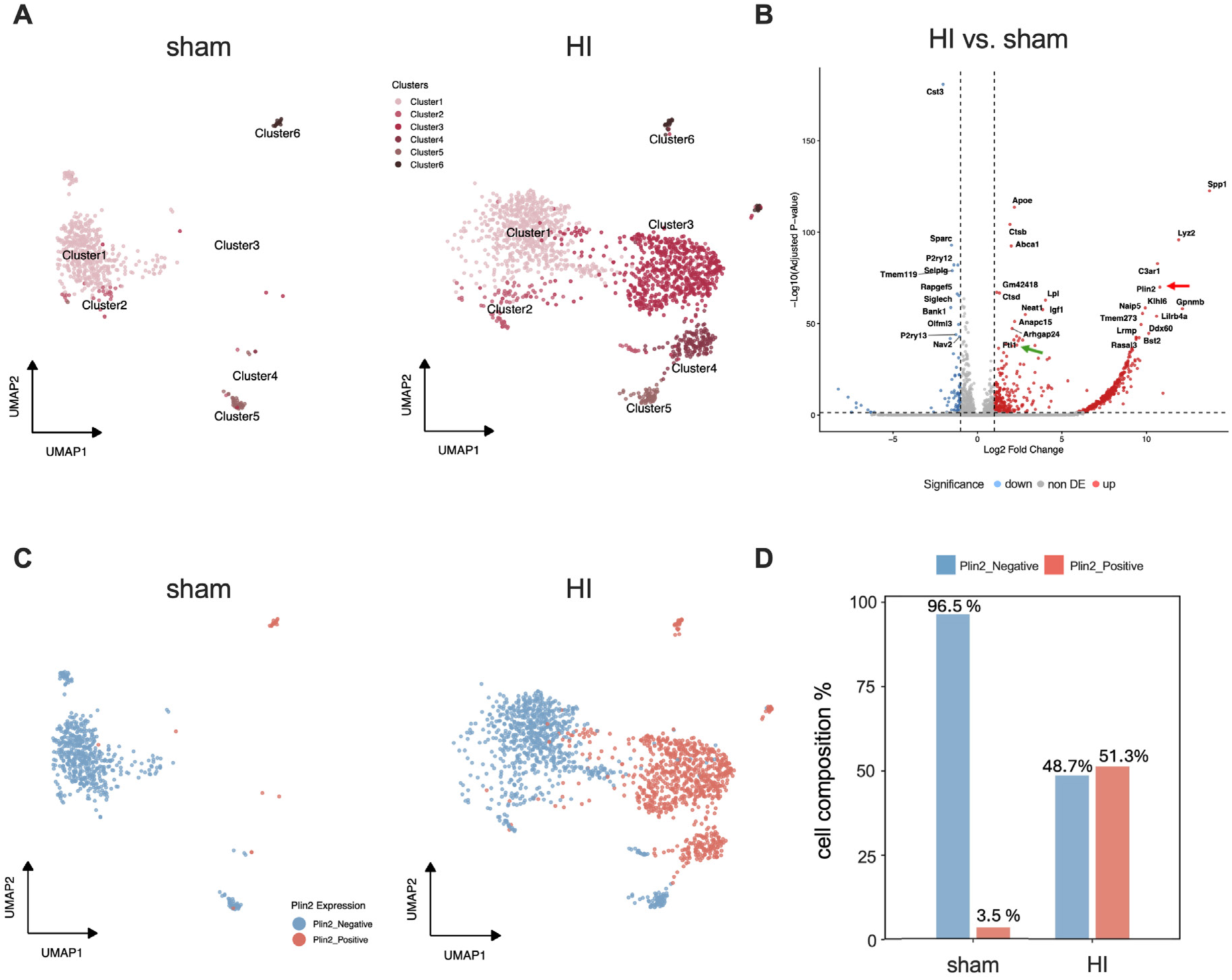
*Plin2*-positive microglial clusters emerge after neonatal HI. A). UMAP projection plot showing the six microglial clusters from the sham and HI-injured ipsilateral cortices at 72hr after HI. New clusters 3 and 4 emerged after HI. n=3 biological replicates for each condition. B). Volcano plot of the top 35 differentially expressed genes in microglia at 72hr after HI as compared to the sham animals. The genes on the right (red) were upregulated, and those on the left (blue) were downregulated. *Plin2* (red arrow) and *Ftl1* (green arrow) genes are marked. C). Cluster reanalysis segregating the *Plin2*-positive (red) and *Plin2*-negative (blue) microglia illustrates that *Plin2*+ cells were mapped into the new clusters 3 and 4. D). The proportion of *Plin2*-positive and *Plin2*-negative microglia in the sham and HI-injured animals.

Differential gene expression analysis (adjusted p-value <0.05, average |log2FC|>1) revealed that, among 14,574 genes detected in microglia, 561 genes were significantly up-regulated, and 81 genes were down-regulated after HI compared to sham controls. The volcano plot (Fig. 8B) showed that *Plin2* (red arrow) was among the top DEGs. Additionally, increased expression of *Ftl1* gene (green arrow), encoding the light chain subunit of the main iron storage protein ferritin, may partially explain iron deposition observed in ischemic regions shown in Fig. 3.

### LD-accumulating microglia (LDAM) have distinct transcriptional signature after HI

To determine whether LDAM exhibit a different transcriptional profile, we re-analyzed microglia snRNA-seq data based on *Plin2* gene expression. Interestingly, *Plin2*-positive cells were mapped into the two newly emerged cluster 3 and cluster 4 (Fig. 8C), suggesting that HI-responsive microglia express *Plin2* and accumulate LD at 72hr after HI. The percentage of Plin2+ microglia over total number of microglia was increased from 3.5% in the sham group to 51.3% in the HI group (Fig. 8D).

Cluster reanalysis segregating *Plin2*-positive from *Plin2*-negative microglia in the HI group revealed marked transcriptomic differences, with 498 genes being upregulated and 157 genes downregulated in the *Plin2+* cells versus *Plin2-*negative counterparts. Among the top 35 up- or down-regulated DEGs (Fig. 9A), expression of homeostatic marker genes, including *P2ry12, Tmem119, Cst3* and *Siglech*, was decreased in *Plin2*+ microglia. Colony-stimulating factor-1 receptor (*Csf1r*), essential for microglial survival, proliferation and differentiation, was also reduced [53, 54]. Conversely, key regulators of microglial activation, such as *Spp1*, *Gpnmb*, and *Lilrb4a*, were significantly upregulated in the *Plin2+* microglia. Elevated expression of *Ctsb, Ctsd* and *Ctsz*, encoding lysosomal proteases, together with the reduced expression of *Cst3* gene, encoding a cystine protease inhibitor, suggests enhanced lysosomal and phagocytic activity in *Plin2+* microglia. Another lysosomal marker for microglia activation, *Cd68* [55, 56], was also increased (log2FC=1.174, adjusted p-value=6.07E-38). These changes suggest that LDAM adopted a less homeostatic and more reactive phenotype.

**Fig. 9:**
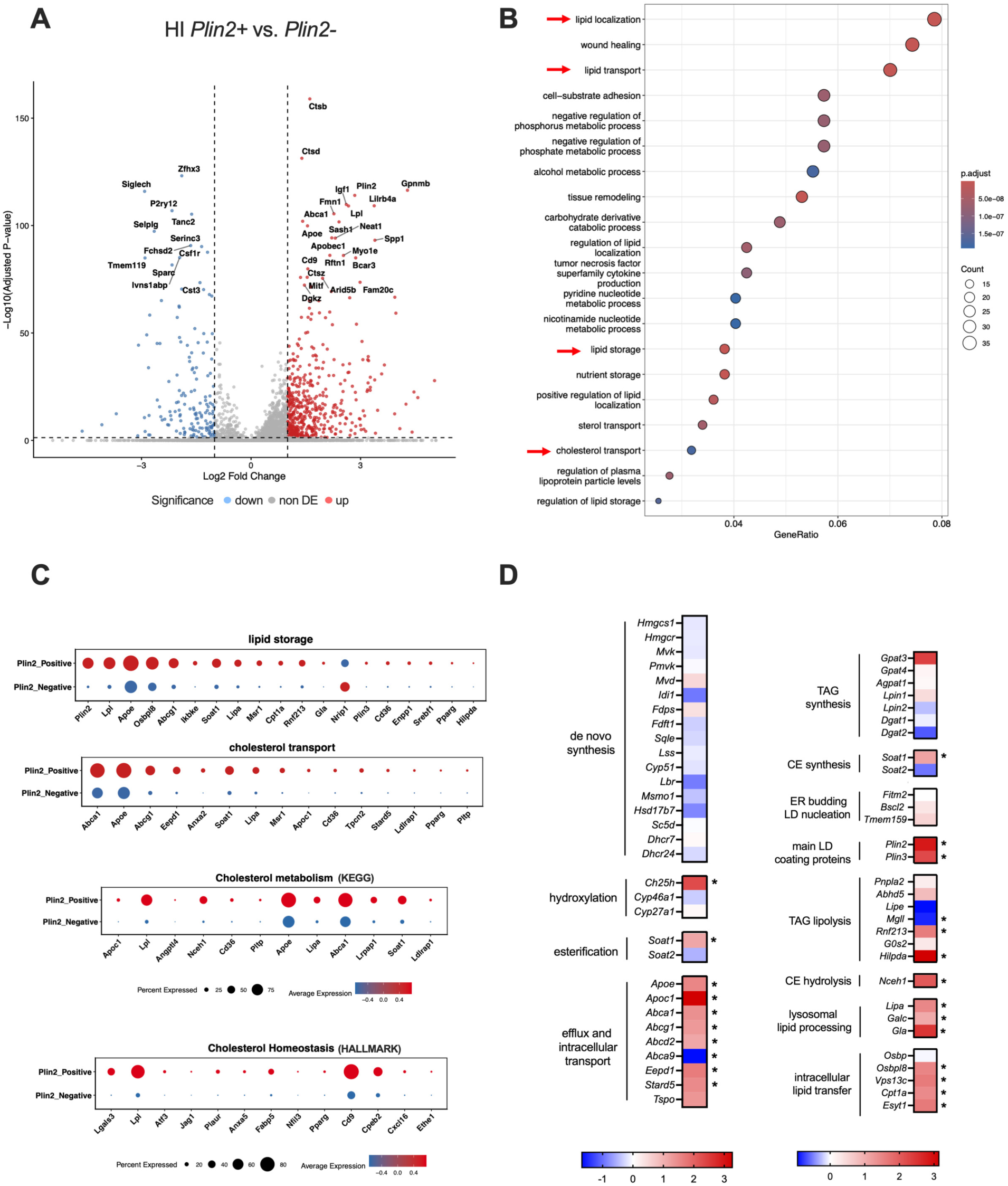
Transcriptional upregulation of genes associated with cholesterol and lipid metabolism in LDAM after HI. A). Volcano plot of the top 35 DEGs in the *Plin2*+ microglia as compared to the *Plin2*-microglia in HI animals. The genes on the right (red) were upregulated, and those on the left (blue) were downregulated. B). Dot plot presenting the significant GO biological processes enriched using upregulated DEGs in *Plin2*+ microglia versus *Plin2*-negative microglia in HI animals. The circle sizes represent the gene counts assigned to particular GO term, and circle colors represent the adjusted p-value significance. The gene ratio on the x-axis represents the proportion of DEGs that are annotated to a specific GO term relative to the total number of genes associated with that term in the reference database. C). Dot plots showing the specific genes annotated in the indicated GO terms, “cholesterol metabolism” pathway using KEGG gene sets, and “cholesterol homeostasis” using Hallmark gene sets. The circle color represents the average expression level, and the circle sizes show the percentage of cells in each group that expressed the gene. D): The heatmap showing the expression levels (log2FC) of the genes involved in cholesterol metabolism (left) and the genes regulating lipid droplet bioprocess (right) in *Plin2*+ LDAM versus *Plin2*-negative microglia. * adjusted p value <0.05.

Genes involved in lipid metabolism, *Apoe, Lpl*, *Abca1* and *Apobec1*, commonly induced in disease-associated microglia (DAM) or injury-responsive states [57, 58], were upregulated in *Plin2*+ LDAM. Both *Ftl1* and *Fth1*, encoding the ferritin light- and heavy-chain subunits, were upregulated in LDAM (log2FC=1.874, Padj=4.37E-60 for *Ftl1*; log2FC=1.119, Padj=2.26E-21 for *Fth1*), suggesting enhanced intracellular iron sequestration consistent with an iron-stressed, activated microglial phenotype.

### Transcriptional upregulation of genes involved in cholesterol and lipid droplet metabolism in LDAM

GO analysis of upregulated DEGs in *Plin2*+ LDAM versus *Plin2*-negative microglia after HI revealed enrichment of several lipid-related biological processes, including lipid localization, lipid transport, lipid storage and cholesterol transport (Fig. 9B). Using the same DEGs also demonstrated enrichment of “cholesterol homeostasis” in the Hallmark dataset and “cholesterol metabolism” in the KEGG database (Fig. 9C). Differences in individual genes within these pathways are shown in Fig. 9C. Given the complexity of lipid species generated following HI and from the breakdown of myelin and apoptotic neurons, we focused our analysis on genes governing cholesterol homeostasis and lipid droplet regulation (Fig. 9D), which included some of the genes listed in these enriched pathways.

#### Genes regulating cholesterol homeostasis

The heatmap (Fig. 9D) illustrates the changes of key genes involved in cholesterol de novo synthesis, hydroxylation, esterification for LD storage, intracellular transport and efflux. Most of the genes encoding enzymes responsible for *de novo* cholesterol synthesis showed a trend of decrease in LDAM, although the changes did not reach statistical differences. In contrast, genes involved in cholesterol hydroxylation (*Ch25h*), cholesterol esterification (*Soat1*), and cholesterol efflux (*Apoe, Apoc1, Abca1, Abcg1, Abcd2, Eepd1* and *Stard5*) were significantly upregulated. To validate the elevated *Ch25h* expression, we measured 25-HC concentrations in the brain. As shown in Fig.10, 25-HC levels in the ipsilateral cortices were markedly higher than those in the sham and the contralateral cortices at 24hr and 72hr post-HI.

**Fig. 10:**
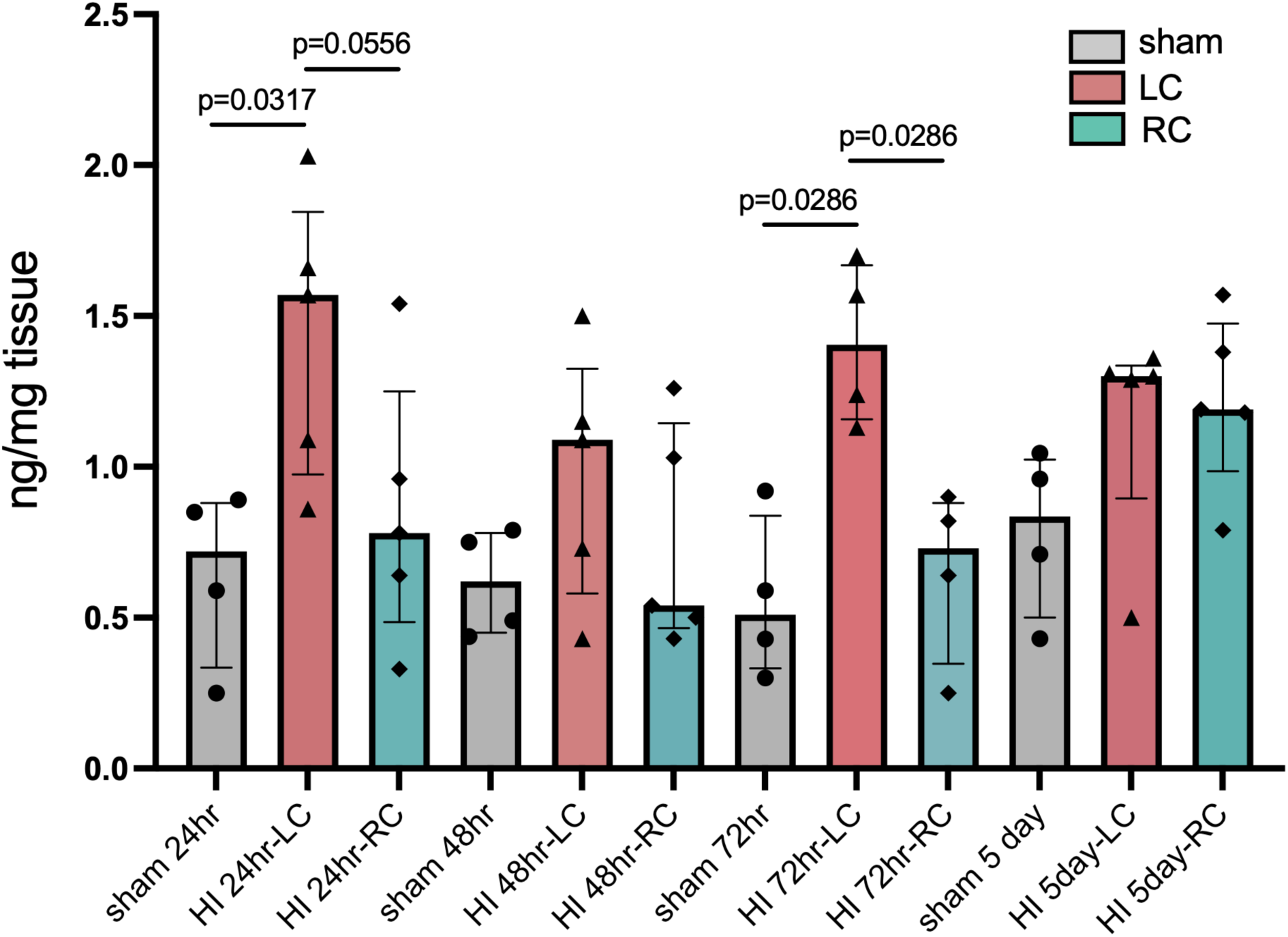
Increased production of 25-HC in the ipsilateral cortices following neonatal HI. The levels of 25-HC were increased at 24hr and 72hr after HI. The amounts of 25-HC were normalized to the tissue wet weight (ng/mg tissue weight). LC: left cortex (ipsilateral side), RC: right cortex (contralateral side), n=4 for sham animals, n=4-5 for HI-injured animals at each time point.

Collectively, these data imply that LDAM pause cholesterol neogenesis while activating pathways that facilitate hydroxylation, export and storage of surplus cholesterol within the cells.

#### Lipid-droplet related genes

In LDAM, lipid accumulation outpaces lysosomal digestion, leading to secondary storage of cholesterol and FA in lipid droplets. We created a heatmap highlighting the changes of important genes regulating LD formation, accumulation and breakdown processes (Fig. 9D). At the transcriptional level, the genes for TAG synthesis remained unchanged, while *Soat1*, catalyzing cholesterol esterification, was upregulated. Increased LD-coating proteins PLIN2 and PLIN3 can stabilize LD, promote their growth and prevent LD lipolysis and autophagy [59, 60]. LD are degraded through two routes: cytosolic lipolysis and lysosomal lipophagy. TAG lipolysis is likely suppressed, as downregulation of *Mgll* (encoding MGL, which breaks down monoglycerides into FFA and glycerol) limits monoglyceride hydrolysis. In parallel, upregulation of *Rnf213*, encoding E3 ubiquitin ligase ring finger protein 213, promotes removal of ATGL from LD surface, thereby reducing TAG breakdown and stabilizing LD [61, 62]. Overexpression of RNF213 in Hela cells or zebrafish increases both the quantity and size of cellular LD [62]. CE hydrolysis may be enhanced as a result of increased expression of *Nceh1* (encoding neutral cholesterol ester hydrolase 1). Enzymes involved in lysosomal lipid processing during lipophagy were upregulated. *Lipa* encodes LAL, crucial for breaking down TAG and CE within lysosomes into free cholesterol and fatty acids [63]. The increased transcription of *Galc* and *Gla* genes suggest an upregulation of lysosomal processing of myelin and membrane-derived lipids during phagocytosis.

The lipid products generated by lysosomal degradation, including FA and cholesterol, can be recycled and redistributed to other organelles. LD constantly form membrane contact sites (MCSs) with other organelles, such as ER, mitochondria, Golgi apparatus, endosomes and peroxisomes, for lipid and protein exchange [64]. We found increased expression of multiple genes in LDAM that are involved in intracellular FA transportation. These include *Fabp5* [65], encoding a member of fatty acid-binding proteins (FABP), the primary “chaperones” to shuttle FA within the cytoplasm to different organelles; *Cpt1a* and *Esyt1* genes, which proteins help transport FA from LD to mitochondria for β oxidation [66, 67]; as well as oxysterol-binding protein-like (Osbpl) family member *Osbpl8*, encoding ORP8 that binds and shuttles oxysterols, cholesterol and phospholipids between membranes [68].

Taken together, the integrated pattern of these alterations in LDAM suggests a strong drive toward LD formation and stabilization, facilitated intracellular lipid trafficking to maintain metabolic balance under phagocytic stress, and increased LD turnover via lipophagy rather than classical lipolysis.

### Transcriptional upregulation of inflammation-associated pathways in LDAM after HI

Lipids and lipid-based signaling molecules are critical regulator of neuroinflammation [69, 70]. We speculated that the distinct lipid gene expression in LDAM modulates their immune responses. Hallmark pathway analysis of upregulated DEGs in LDAM demonstrated enrichment of multiple inflammation-associated pathways (Fig. 11A). These included TNFα signaling via NF-κB, interferon-γ response, Il2-Stat5 signaling, and Il6-Jak-Stat3 signaling. The “Inflammatory response” pathway was enriched with the specific genes listed in Fig. 11C. A number of upregulated genes in these pathways (Fig. 11C), such as *Abca1, Lpl, Msr1, Cd36, Plaur* and *Thbs1*, are regulators of lipid metabolism, suggesting a connection between altered lipid molecules in LDAM and their potentially different inflammatory responses compared to their LD-free counterparts.

**Fig. 11:**
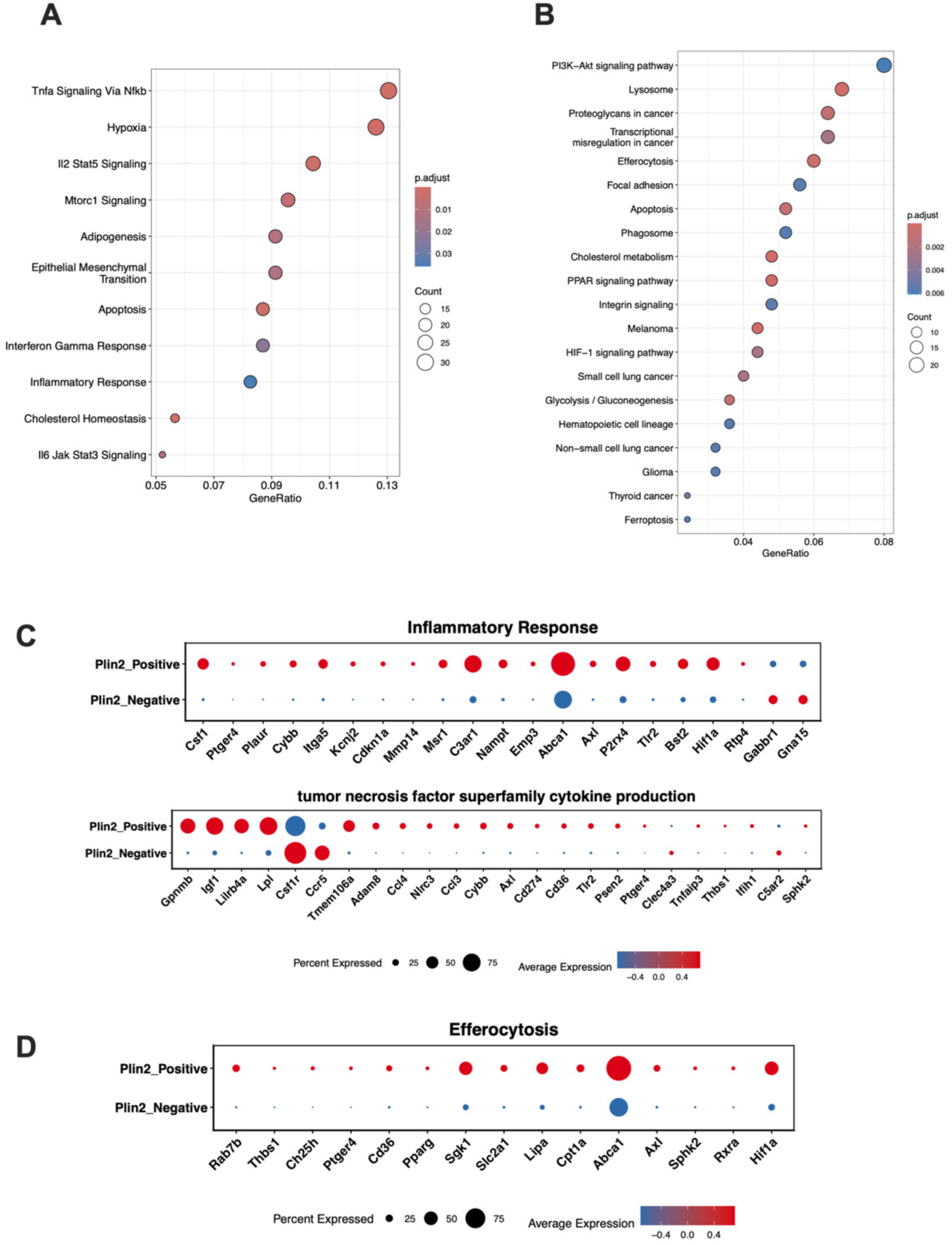
Upregulated pathways and associated genes in *Plin2*+ LDAM vs. *Plin2*-negative microglia after HI. Dot plots showing the pathways enriched with upregulated DEGs in *Plin2*+ LDAM vs. *Plin2*-negative microglia after HI using Hallmark gene sets (A) or KEGG gene sets (B). C) and D): Dot plots presenting the specific genes annotated in the indicated pathways. The circle color represents the average expression level, and the circle sizes show the percentage of cells in each group that expressed the gene.

### Genes in the phagocytosis pathways are upregulated in LDAM following HI

At the transcriptional level, LDAM appear to have increased phagocytosis. The “efferocytosis”, “phagosome” and “lysosome” pathways were enriched in KEGG analyses (Fig. 11B), and many genes in the efferocytosis pathway are related to lipid metabolism (*Abca1, Lipa, Cpt1a, Ch25h, Pparg, Sphk2*, Fig. 11D). Expression of *Lgals3*, encoding galectin-3 that is a marker of activated/phagocytic microglia [71, 72], was increased in LDAM (Fig. 9C), along with genes for the phagocytic receptors *Axl* and *Cd36*. ABCA1 promotes macrophage efferocytosis in the context of atherosclerosis [73], beyond its classic function in cholesterol efflux. In an AD mouse model, increased ABCA1 is associated with improved microglial Aβ phagocytosis[74]. Upregulation of ABCA1 is also required for the transformation of reactive astrocytes into a phagocytic phenotype after transient brain ischemia [75].

To validate whether LD accumulation is associated with enhanced phagocytosis and inflammatory responses after HI, we treated mouse microglia-derived BV2 cells in the present of DGAT inhibitors. DGAT1/2 are the key enzymes in the final, committed step of TAG biosynthesis (Fig.1) [76]. We found a significant increase in the percentage and expression of both PLIN2+ (Fig. 12A, 12B) and BODIPY+ (Fig. 13A, 13B) cells after OGD. Treatment with either the DGAT1 (T863) or DGAT2 inhibitor (PF) markedly reduced OGD-induced LD accumulation (Fig.12, 13). To study whether LD-containing microglia had altered phagocytic activity, we incubated BV2 cells with FITC-dextran 40 after OGD (Fig. 12A, 12E-G). OGD increased microglia phagocytosis with nearly all BV2 cells in the blank (medium) and DMSO group are FITC-dextran-positive. Both inhibitors reduced the percentage of dextran+ cells (Fig.12F) and Dextran 40 fluorescent intensity (Fig. 12G), suggesting LD formation is associated with enhanced phagocytic capacity of BV2 microglial cells following OGD. Interestingly, OGD increased the number and level of BV2 cells expressing IL-1β, while inhibition of DGAT-mediated LD biogenesis diminished the proportion of IL-1β-positive BV2 cells (Fig. 13F) and IL-1β fluorescent intensity (Fig. 13G).

**Fig. 12:**
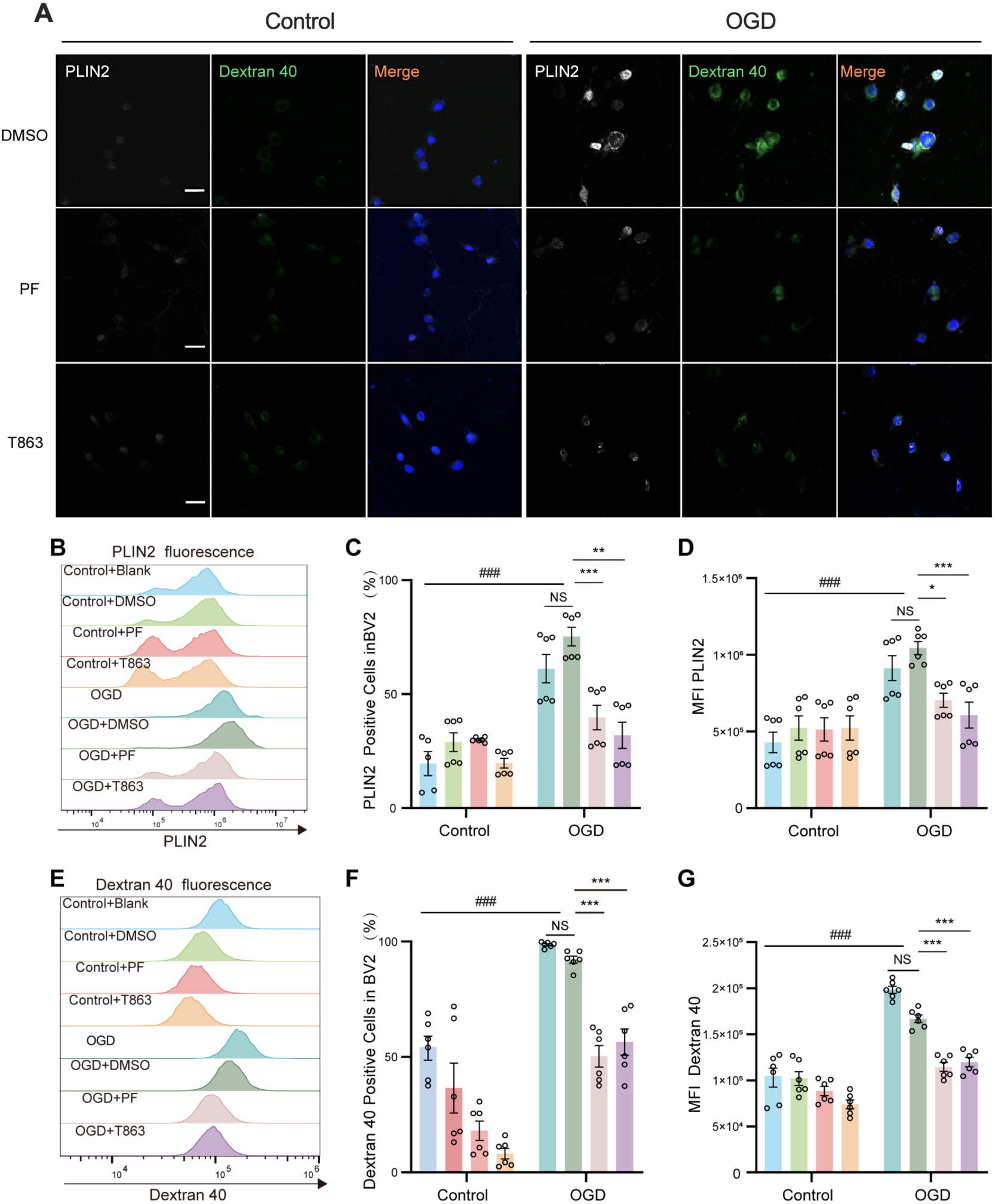
DGAT inhibitors attenuate OGD-induced increase in PLIN2 expression and phagocytic activity in microglia. A). Representative images of PLIN2 (white) expression and Dextran uptake (green) in BV2 cells following DGAT inhibitor treatment. BV2 cells were treated with DGAT1 inhibitor T863 or DGAT2 inhibitor PF-06424439 (PF) or DMSO vehicle control under normoxic control or OGD condition. Scale bar = 20 µm. n = 3 per group. B). Representative flow cytometric histograms showing PLIN2 fluorescence intensity curves under different treatment conditions. Blank group cells received medium as no-treatment control. C). Percentage of PLIN2^+^ cells in BV2 cells under different treatment conditions. D). PLIN2 mean fluorescence intensity (MFI) in BV2 cells under different treatment conditions. E). Representative flow cytometric histograms showing Dextran 40 fluorescence intensity curves under different conditions. F). Percentage of Dextran 40^+^ cells in BV2 cells under different treatment conditions. G). Dextran 40 MFI under different treatment conditions. For flow cytometry, n = 6 per group; data are presented as mean ± SD; two-way ANOVA (OGD × drug) with Tukey’s multiple comparisons test; *p < 0.05, **p < 0.01, ***p < 0.001 vs. OGD and DMSO group; ### p < 0.001 vs. OGD + Blank group.

**Fig. 13:**
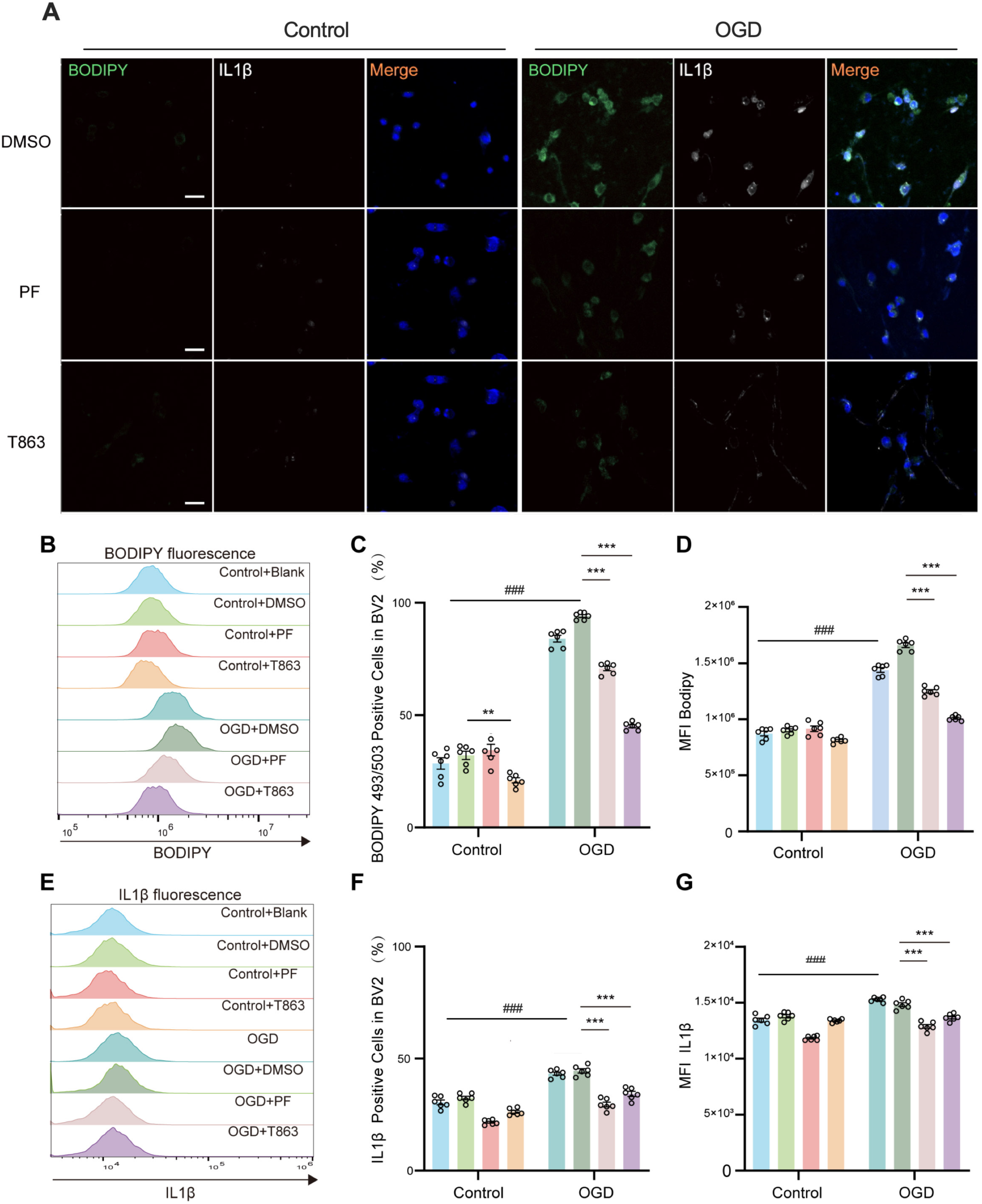
DGAT inhibitors attenuate OGD-induced lipid droplet accumulation and IL-1β expression in microglia. A). Representative images of BODIPY (green) and IL-1β (white) expression in BV2 cells following DGAT inhibitor treatment. BV2 cells were treated with DGAT1 inhibitor T863 or DGAT2 inhibitor PF-06424439 (PF) or DMSO vehicle control under normoxic control or OGD condition. Scale bar = 20 µm. n = 3 per group. B). Representative flow cytometric histograms showing BODIPY fluorescence intensity curves in BV2 cells under different treatment conditions. Blank group cells received medium as no-treatment control. C). Percentage of BODIPY^+^ cells in BV2 cells under different treatment conditions. D). BODIPY mean fluorescence intensity (MFI) under different treatment conditions. E). Representative flow cytometric histograms showing IL-1β fluorescence intensity curves under different treatment conditions. F). Percentage of IL-1β^+^ cells in BV2 cells under different treatment conditions. G). IL-1β MFI in BV2 cells under different treatment conditions. For flow cytometry, n = 6 per group; data are presented as mean ± SD; two-way ANOVA (OGD × drug) with Tukey’s multiple comparisons test; *p < 0.05, **p < 0.01, ***p < 0.001 vs. OGD and DMSO group; ### p < 0.001 vs. OGD + Blank.

In vivo, at 72hr after HI, most PLIN2+ microglia expressed IL-6 (∼ 70%) and IL-1β (∼ 80%) with a significantly higher expressing level than PLIN2-microglia (Fig. 6D, 6E) suggesting a correlation between lipid droplet accumulation and pro-inflammatory cytokine expression. Together, the in vivo and in vitro results support a role for lipid droplets in promoting microglial inflammatory responses and phagocytic activities.

## Discussion

In this study, we demonstrated that lipid droplets build up predominately in microglia after HI in injured brain regions in neonatal mice, and in human HIE brain as well. At transcriptional level, LD accumulation represents an alteration in microglial metabolism, where LDAM display an upregulation of genes associated with microglial activation, enhanced cholesterol and lipid processing, phagocytic and pro-inflammatory phenotype. In neonatal mice, LDAM is associated with increased expression of cytokines IL-6 and IL-1β after HI. In BV2 microglial culture, pharmacological inhibition of LD biogenesis attenuates OGD-induced lipid droplet accumulation, phagocytic activity, and IL-1β expression. These results suggest that lipid droplet formation is functionally linked to microglia-mediated phagocytic and inflammatory responses following HI.

This is the first characterization of lipid droplet-accumulating cells after neonatal brain hypoxia-ischemia. We reported the changing dynamics of PLIN2-positve microglia and infiltrating monocyte-derived macrophages (MDM) in the first week after HI. Although both microglia and MDM accumulate LD, the majority of LD-containing myeloid cells are resident microglia rather than infiltrating monocytes. Lipid droplet accumulation peaks at 3 days after injury and persists through at least 7 days in microglia. In contrast, Plin2+ MDM decrease over time. The results from P2ry12CreER;Ai14 mice, a highly specific microglia-reporter line, reaffirmed that LD are mainly localized in microglia. This is in line with a recent report in adult stroke model demonstrating that the principal stroke-associated myeloid lineage cells are resident microglia with a lipid-phagocytosing and increased lipid clearance phenotype [17]. LD were not detected in neurons or astrocytes, consistent with findings from adult stroke model showing minimal lipid droplets accumulation in these cell types during the acute phase of stroke [77]. Reactive phagocytic astrocytes have been reported following adult brain ischemia, but they appear at a later stage (peaked 7 days post-HI) at penumbra region, while phagocytic microglia are located within the ischemic core at an earlier time (peaked 3 days) [75], similar to our findings. We have also shown that at 72hr after neonatal HI, microglia are the major phagocytes [9]. A previous study in the Vannucci neonatal HI model reported that MDM infiltration peaks early and declines significantly by day 7, whereas microglial numbers peak at day 3 and remain elevated through day 7 [78]. We did not include the total numbers of microglia and MDM in this study, but the changes in lipid droplet-accumulating microglia or MDM followed the same pattern (Fig. 6B, 6C) indicating a possible temporal association between lipid droplet formation and myeloid cell responses after HI.

Recent studies suggest that MDM undergo progressive reprogramming and acquire a microglia-like phenotype in the ischemic brain [79, 80]. For example, recruited monocytes frequently down-regulate CCR2 expression while concomitantly upregulating CX3CR1, representing a dynamic recruitment-to-adaptation-to-residency of myeloid states [81]. This phenotypic convergence following brain injury makes the two populations difficult to distinguish with current markers. Additional markers and advanced fate-mapping approaches could help elucidate shared and distinct metabolic programs in MDM and microglia at early-stage inflammatory induction and late-stage inflammatory resolution during brain recovery and repair. Lipid droplets accumulation in MDM has been reported after ischemic stroke in adult animals [82–84]. A recent study investigating Ly6C^high^ Ly6G^low^ MDM in the ischemic brain showed that infiltrating MDM undergo lipid metabolic reprogramming in the stroke brain. HIF1α-regulated hypoxia-inducible lipid droplet associated protein (HILPDA) was identified as a key regulator of their adaptation to the ischemic environment, promoting phosphatidylcholine synthesis and an anti-inflammatory phenotype [83]. Both HIF1α signaling pathway and *Hilpda* gene was found upregulated in *Plin2+* microglia in our snRNA-seq data (Fig. 11B, Fig.9C), although the percentage of *Plin2+* microglia expressing *Hilpda* gene was very low (Fig. 9C). It would be interesting to profile *Plin2+* MDM transcriptome at 24 hour after HI to identify their phenotype and roles in early inflammatory responses. Given that microglia are the major lipid droplet-accumulating cells in our HI model, this study focused primarily on their role in the post-HI response. In snRNA-seq data analyses, we used a combination of four specific genes (*P2ry12*, *Cx3cr1, Itgam* and *C1qa*) to identify microglia, which is rigorous.

LD formation is a microglial response to HI, as 51.3% of microglia became *Plin2*-positive after HI compared with only 3.5% in the sham animals. Importantly, *Plin2*-expressing cells clustered within the newly emerged HI-responsive microglia subclusters. The transcriptional pattern of LDAM suggest a transition from homeostatic to glycolytic immune-active state, which may support phagocytosis and debris clearance, especially in early injury responses. The upregulated lipid-related genes in LDAM indicates accelerated cholesterol and lipid handling, including 1). increased lipid uptake and storage rather than cholesterol neogenesis; 2). enhanced lipid transfer at membrane contact sites, highlighting close functional coupling between LD and other organelles; 3). LD turnover rely more heavily on lipophagy that is slower but more organized than cytosolic lipolysis. Reduced lipolysis favors LD accumulation. 4). enhanced lipid clearance including hydroxylation and export of cholesterol and oxysterols. Together, these changes imply a coordinated effort to maintain proper lipid balance and cellular function. However, microglia may be overwhelmed by persistent LD accumulation and become dysfunctional, contributing to a delayed resolution of inflammation and brain recovery.

Recent studies suggest that triglyceride accumulation and LD biogenesis are necessary for LPS-mediated activation of human iPSC-derived microglia, as well as primary mouse microglia [85]. Activated microglia have high energetic demands, and LD could act as metabolic hubs supporting activated microglia. Manipulating TAG metabolism and associated-LD abundancy is also linked to changes in LPS-induced microglial phagocytosis and related inflammatory and behavioral responses in vivo [86], demonstrating a role for LD in microglia activation and brain inflammation. In our study, LDAM exhibited transcriptional upregulation of multiple inflammation-associated pathways, suggesting a shift towards pro-inflammatory phenotype. These changes were supported by the in vivo and in vitro data showing that PLIN2+ microglia had increased proportion and higher expression of IL-6 and IL-1β compared to PLIN2-microglia at 72hr post-HI. In BV2 cells, reducing lipid droplets formation attenuated IL-1β expression. Similar results were reported in primary microglia showing association of lipid droplet-rich microglia with elevated IL-1β, IL-6, and TNFα production following OGD treatment [87]. These findings support a role for LD in the modulation of HI-induced neuroinflammation. The underlying mechanisms are not completely understood. Lipid droplets may provide substrates for inflammatory lipid mediators as they store TAG and CE that can be mobilized by lipases [88]. Prior work has implicated LD-associated proteins, for example PLIN2, in neuroinflammation. PLIN2 knockdown has shown to block NLRP3 inflammasome activation and attenuate cerebral ischemia/reperfusion injury in adult SD rats [89]. Moreover, PLIN2 expression is elevated in patients with early tauopathy or AD and is associated with Il-6 expression [90]. Microglial maturation stage, diversity and plasticity are critical in their responses to acute CNS injuries. A recent study using fate mapping of *Spp1*-expressing DAM like cells after stroke revealed age-dependent plasticity, where neonatal (P6) DAM-like cells can regain homeostasis as stroke resolves, while in juvenile mice (P28), these cells are irreversibly removed after stroke [91]. Early postnatal microglia have high heterogeneity and plasticity compared to adult microglia [92–94]. It would be valuable to trace LDAM longitudinally in our model using a LD-reporter mouse line, for example a fluorescent Plin2 knock-in mouse strain [95]. In summary, this is the first study of microglial lipid droplets using a clinically relevant HIE mouse model. We demonstrate that lipid droplets-accumulating microglia displayed a distinct immuno-metabolic transcriptional landscape. At 72 hours or early phase after HI, acute lipid droplet accumulation may represent a microglial activation state associated with enhanced phagocytosis and increased pro-inflammatory cytokine production. This differs from LDAM in the aging brain, where microglia undergo chronic and persistent lipid droplet accumulation leading to an impaired and dysfunctional phenotype [33]. Both protective and detrimental role of LDAM in adult stroke models have been reported [32, 77, 82, 96]. As PLIN2-positive LDAM still exist at 7 days after HI (∼30% of total microglia in the ipsilateral cortex, Fig. 6B), characterization of this microglia subset at later time points can help us further understand how lipid processing controls microglial functional states; and contributes to HI brain injury or regeneration. LDAM in our animal model closely resemble those in HIE human brains highlighting a translational potential of the work.

Several key questions remain unanswered. Can LDAM in the developing brain regain homeostasis or are they primed for elimination, and by which mechanisms? Does LD accumulation contribute to prolonged microglial activation and chronic inflammation and hinder HI recovery [97] or do they support repair? Could interventions abrogating LDAM at subacute stage mitigate neonatal HI brain injury and improve long-term outcomes? Future work should address these critical questions.

## Declarations

### Ethics approval and consent to participate

Brain sections from PFA-fixed, OCT-embedded human HIE brain tissue were used in this study. De-identified human tissue was collected from the Autopsy Service in the Department of Pathology at UCSF, with previous patient consent in strict observance of the legal and institutional ethical regulations.

All animal procedures were performed according to the guidelines of the Institutional Animal Care and Use Committee at UCSF and Shantou University Medical College.

### Consent for publication

Not applicable

### Availability of data and materials

The snRNA-seq data generated and/or analyzed during the current study are available in the Open Archive for Miscellaneous Data (https://ngdc.cncb.ac.cn/omix/: accession no: OMIX014659). All other data are available from the corresponding authors on reasonable request.

### Author’s contributions

Lei, S. Li: acquisition and analysis of data; G. Zhang, Y. Li: acquisition of data; B. Wu: acquisition of data; P Pan: acquisition of data; D. Ferriero and Z. Guan: revising article critically for important intellectual content; F. Li: guidance for in vivo and in vitro flow cytometry experiments and data analysis; F. Li and X. Jiang: substantial contributions to conception and design, interpretation of data, drafting the article and revising article critically for important intellectual content, and approving the version to be published.

## Figure Legend

**Supplementary Fig. 1: Lipid droplets are not present in neurons and astrocytes after HI in neonatal mice** A): PLIN2 (green) was expressed in the ipsilateral (ipsi-), but not in the contralateral (contra-) cortex. The neurons in the ipsilateral cortex were shrunken compared to the healthy neurons in the contralateral cortex. There was no colocalization of PLIN2 (green) with NeuN staining (blue). B): Reactive astrocytes in the ipsilateral cortex had pronounced hypertrophy of the cell body and processes compared to the healthy astrocytes in the contralateral cortex. PLIN2 (green) was not expressed in GFAP (blue)-positive astrocytes (n=3).

**Supplementary Fig. 2: Human neurons and astrocytes do not accumulate lipid droplets in HIE brain** The brain sections from human HIE cases in Fig. 7 were double stained with PLIN2 antibody (red) paired with anti-GFAP (A: green) or anti-NeuN (B: green) antibodies. PLIN2 expression was not observed in neurons and astrocytes.

**Supplementary Fig. 3: Identification of brain cell types via snRNA-seq analysis and annotation** A): UMAP projection plot presenting identified cell clusters harvested from the ipsilateral brain at 72hr after HI B): UMAP plots showing expression of the four selected microglial marker genes in brain cell clusters C): Dot plot of the top four marker genes for each cell type; dot size represents the percentage of cells expressing the gene, and color indicates scaled average expression levels.

**Supplementary Fig. 4: Gene expression heatmap defining six transcriptionally distinct microglia** Heatmap showing scaled expression (z-score) of representative marker genes across six clusters (cluster 1-6). Columns represent individual cells, grouped by clusters as indicated by the top annotation bar, and rows represent selected marker genes enriched in each cluster. Color intensity indicates relative gene expression, with red denoting higher and blue lower expression. For each cluster, the top 10 characteristic marker genes are listed alongside the heatmap.

## Abbreviations

25-HC: 25-hydroxycholesterol
*Abca1*: ATP-binding cassette subfamily A member 1
*Abcd2*: ATP-binding cassette subfamily D member 2
*Abcg1*: ATP-binding cassette subfamily G member 1
*Aldoa*: aldolase, fructose-bisphosphate A
*Apobec1*: Apolipoprotein B mRNA editing enzyme catalytic subunit 1
*Apoc1*: apolipoprotein C1
*Apoe*: apolipoprotein E
*Atg7*: autophagy Related 7
ATGL: Adipose Triglyceride Lipase
*Axl*: Axl receptor tyrosine kinase
BHFDR: Benjamini-Hochberg false discovery rate
BP: Biological Process
CC: Cellular Component
CCA: common carotid artery
CCR2: C-C motif chemokine receptor 2
CE: Cholesterol esters
*C1qa*: complement C1q A chain
CH25H: cholesterol 25-hydroxylase
*Ch25h*: cholesterol 25-hydroxylase
Cpt1a: Carnitine Palmitoyltransferase 1A
*Csf1r*: Colony-stimulating factor-1 receptor
*Cst3*: Cystatin 3
*Ctsb*: Cathepsin B
*Ctsd*: Cathepsin D
*Ctsz*: Cathepsin Z
CV: cresyl violet
*CX3CR1*: C-X3-C motif chemokine receptor 1
DEGs: differentially expressed genes
*Eepd1*: Endonuclease/Exonuclease/Phosphatase Family Domain Containing 1
*Esyt1*: Extended Synaptotagmin-1
*Fabp5*: Fatty Acid Binding Protein 5
FABP: fatty acid-binding proteins
FC: free cholesterol
FC: fold change
FFA: free fatty acids
*Fth1*: ferritin heavy chain 1
*Ftl1*: ferritin light polypeptide 1
*Galc*: galactosylceramidase
GFP: green fluorescent protein
*Gla*: galactosidase alpha
GLUT1: glucose transporter protein type 1
*Gpi1*: glucose-6-phosphate isomerase 1
*Gpnmb*: Glycoprotein Non-Metastatic Melanoma Protein B
GO: Gene Ontology
HI: Hypoxia-ischemia
HIE: Hypoxic-ischemic encephalopathy
HIF-1: hypoxia-inducible factor 1
HSL: Hormone-Sensitive Lipase
HVGs: highly variable genes
*Itgam*: integrin subunit alpha M
LD: Lipid droplets
LDAM: LD-accumulating microglia
*Ldha*: lactate dehydrogenase A
*Lgals3*: lectin, galactoside-binding, soluble, 3
*Lilrb4*: Leukocyte Immunoglobulin Like Receptor B4
*Lipa*: lysosomal acid lipase
*Lpl*: lipoprotein lipase
MCSs: membrane contact sites
MCT4: monocarboxylate transporter 4
MDM: monocyte-derived macrophages
MF: Molecular Function
MGL: Monoglyceride Lipase
*Msr1*: Macrophage Scavenger Receptor 1
*Nceh1*: neutral cholesterol ester hydrolase
OD: optical densities
*Optn*: optineurin
ORO: Oil Red O
*Osbpl*: oxysterol-binding protein-like
*P2ry12*: purinergic receptor P2Y, G-protein coupled, 12
Padj: adjusted p-value
PCs: principal components
PCA: principal component analysis
PFA: paraformaldehyde
*Pfkl*: phosphofructokinase, liver type
*Pfkfb3*: 6-phosphofructo-2-kinase/fructose-2,6-biphosphatase 3
*Pgm1*: phosphoglucomutase 1
*Pkm*: pyruvate kinase M1/2
*Plaur*: plasminogen activator, urokinase receptor
Plin2: perilipin 2
*Pparg*: peroxisome proliferator-activated receptor gamma
PPARγ: peroxisome proliferator-activated receptor gamma
*Rab7b*: RAB7B, member RAS oncogene family
RFP: red fluorescent protein
RIPA buffer: Radioimmunoprecipitation assay buffer
*Rnf213*: E3 ubiquitin ligase ring finger protein 213
*Sgk1*: Serum/Glucocorticoid Regulated Kinase 1
*Siglech*: Sialic acid binding Ig-like lectin H
*Slc2a1*: Solute Carrier Family 2 Member 1
*Slc16a3*: solute carrier family 16 member 3
snRNA-seq: Single-nucleus RNA sequencing
*Sphk2*: sphingosine kinase 2
SOAT1: Sterol O-acyltransferase 1
*Spp1*: Secreted Phosphoprotein 1
*Srebf1*: Sterol Regulatory Element Binding Transcription Factor 1
SREBP1: Sterol regulatory element-binding protein 1
*Stard5*: StAR related lipid transfer domain containing 5
TAG: triacylglycerol
*Thbs1*: Thrombospondin 1
*Tmem119*: Transmembrane Protein 119
*Tpi1*: triosephosphate isomerase 1
T-SNE: t-distributed stochastic neighbour embedding
UMAP: uniform manifold approximation and projection

