## Supplementary figures and images for "Accumulation of Lipid Droplets in Microglia following Neonatal Brain Hypoxia-Ischemia"

### Supplemental figures

**Supplementary Figure 1**

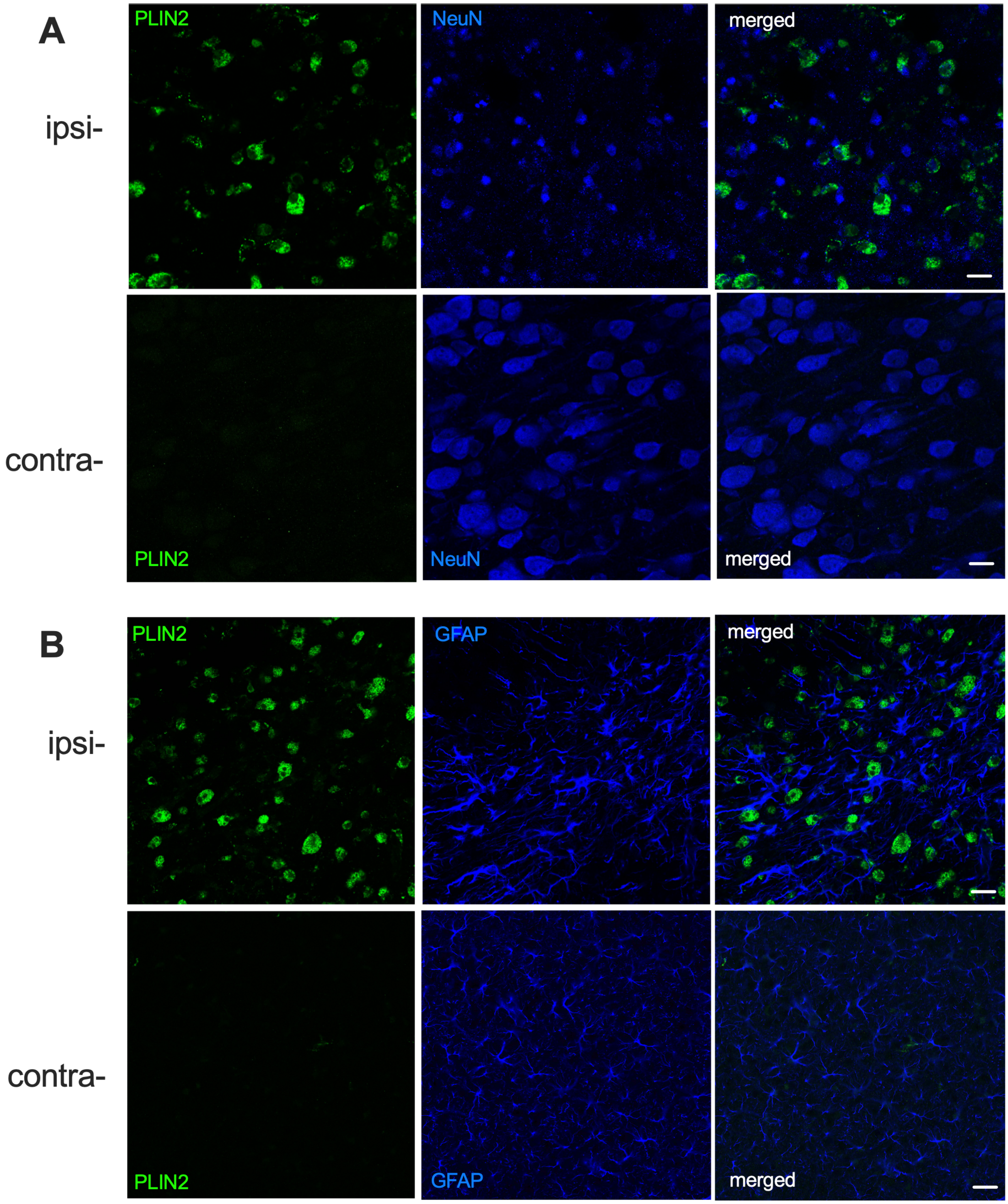

**Supplementary Figure 2**

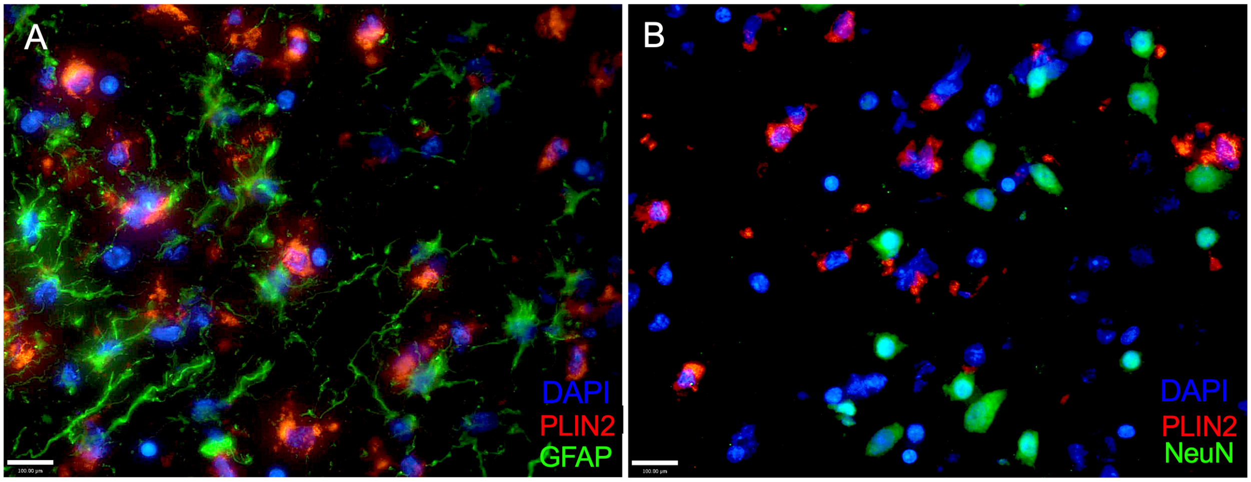

#### Supplementary Figure 3

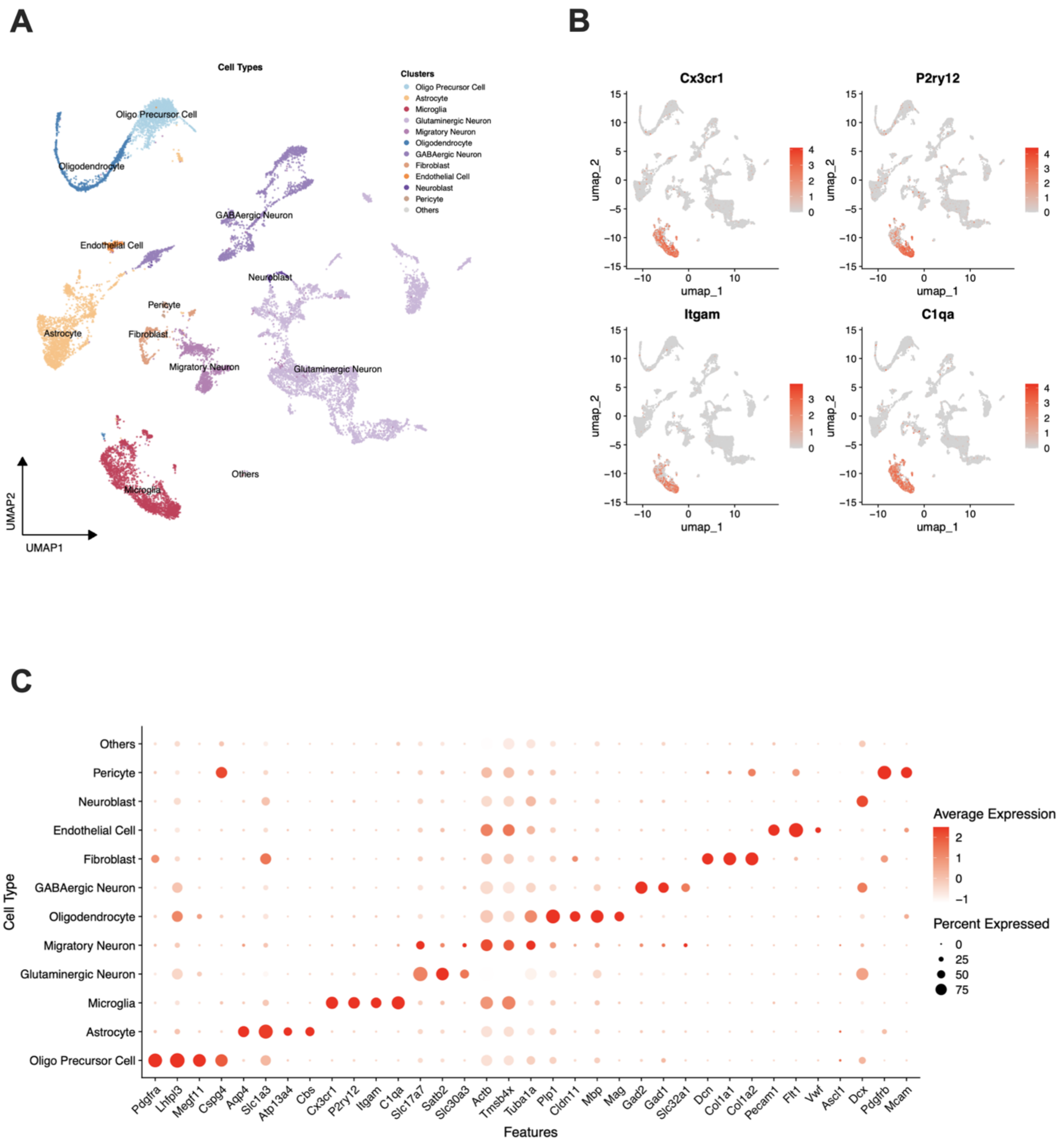

#### Supplementary Figure 4

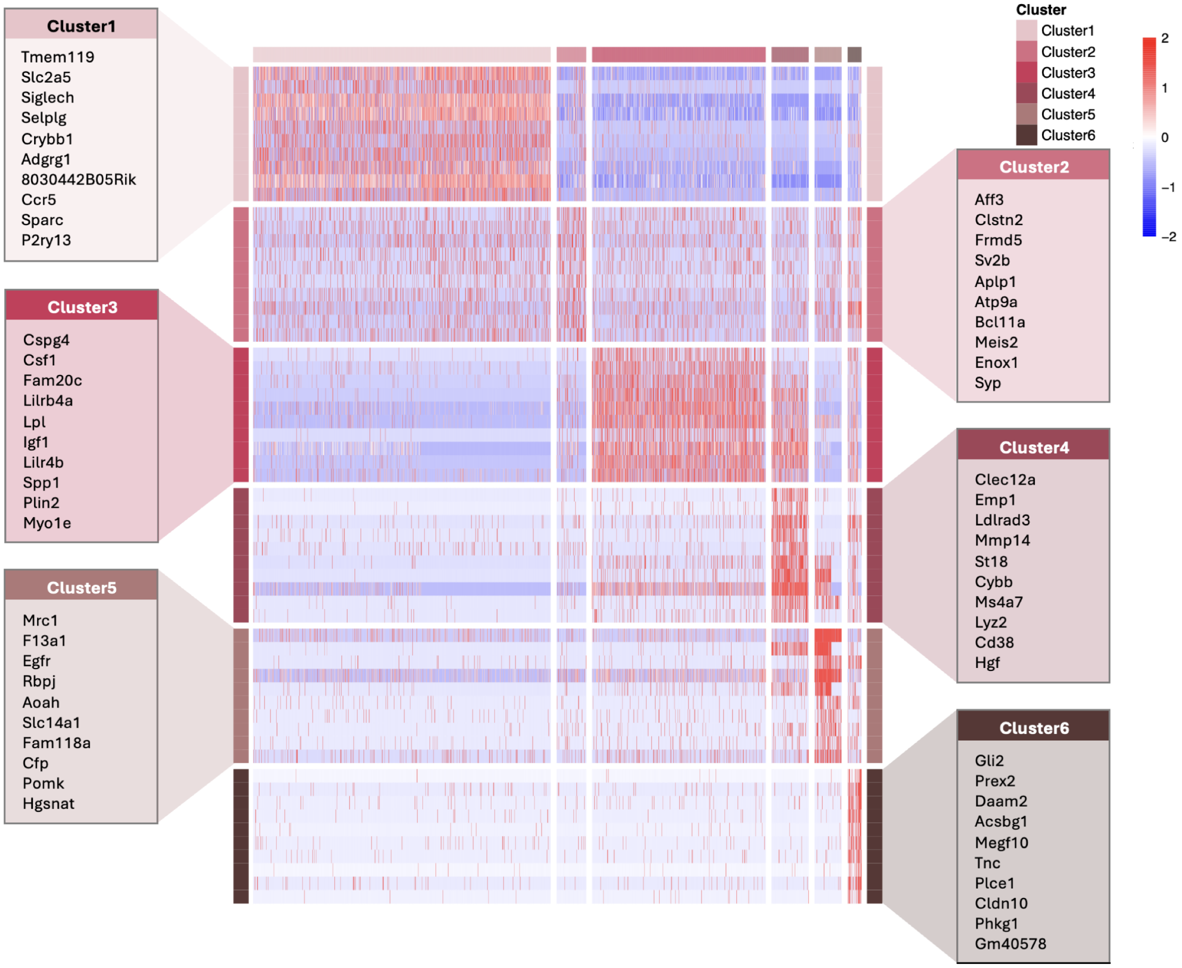
